# Rational inattention in mice

**DOI:** 10.1101/2021.05.26.445807

**Authors:** Nikola Grujic, Jeroen Brus, Denis Burdakov, Rafael Polania

## Abstract

Behavior exhibited by humans and other organisms is generally inconsistent and biased, and thus is often labeled irrational. However, the origins of this seemingly suboptimal behavior remain elusive. We developed a behavioral task and normative framework to reveal how organisms should allocate their limited processing resources such that there is an advantage to being imprecise and biased for a given metabolic investment that guarantees maximal utility. We found that mice act as rational-inattentive agents by adaptively allocating their sensory resources in a way that maximizes reward consumption in novel stimulus-reward association environments. Surprisingly, perception to commonly occurring stimuli was relatively imprecise, however this apparent statistical fallacy implies “awareness” and efficient adaptation to their neurocognitive limitations. Interestingly, distributional reinforcement learning mechanisms efficiently regulate sensory precision via top-down normalization. These findings establish a neurobehavioral foundation for how organisms efficiently perceive and adapt to environmental states of the world within the constraints imposed by neurobiology.

---

Seemingly irrational behavior is surprisingly common in humans and other animals. This has been widely documented in research conducted by neurobiologists, psychologists, and economists, for whom apparent anomalies in the behavior of healthy organisms are difficult to reconcile with idealized statistical and neurobiological frameworks (1, 2). These idealized concepts and theories are increasingly used to guide diagnoses and treatments in the medical domain (3) and policy making in applied economic settings (4, 5), so such anomalies raise an important question: Why are common behavioral strategies so often different from the idealized predictions, and is it reasonable to dismiss such strategies as irrational or suboptimal?

A potential answer to this question might be rooted in a premise that holds across all living organisms: organisms have a limited metabolic budget for interacting with their environments, and thus a restricted capacity to process environmental and interoceptive signals (6). This entails that, for instance, when an organism must decide between various alternatives that promise some reward, the process of choosing the best alternative must be based on imprecise and biased perceptions (7–9). However, it remains poorly understood how organisms can make the best out of such perceptual limitations such that reward is maximized in environments that are uncertain.

One attempt to address this question proposes that in the course of evolution, the nervous system developed computational strategies that take into consideration the statistical structure of the environment and the uncertainty of sensory signals. For instance, theoretical (10), behavioral (11–13) and recent neurophysiological (14, 15) studies support the idea that the brain learns a statistical model of the world and optimally combines this knowledge with imperfect sensory information, and thus approaches optimal computations under uncertainty. However, this approach does not take into consideration the biological limitations of information processing in the nervous system.

This problem has received considerable attention in recent years, when the principles of efficient computation under uncertainty and cognitive limitations can be studied in the rational inattention or resource-rationality framework (16, 17). This approach has been instrumental in accounting for various aspects of behavior that were not possible to explain with rational models not only in decisions relying on basic sensory perception (18, 19) and memory (20, 21), but also in higher-level processes in humans such as the evolution of language (22) and economic decision-making (23–30).

Rational inattention theory predicts that (i) limited time (31, 32), (ii) noise in the system and environment (33, 34), and (iii) metabolic constraints (35) all limit access to information. Consequently, a decision maker may often benefit from violating the axioms of rational modeling frameworks (36–39) by being rationally myopic and ignoring information that is not worth the effort of processing (26, 27, 30, 40). These apparently suboptimal strategies may ultimately lead organisms to maximize their chances of survival. (41, 42).

Nonetheless, formulations of rational inattention, which have to date been applied predominantly in economic settings, often do not display close conformity with neurobiological implementations. Moreover, in some cases these formulations may not be directly translatable to applications of optimal allocation of attentional resources in sensory perception (43). Additionally, previous formulations leave unclear how organisms efficiently adapt and exploit novel stimulus-reward associations when sensory information varies in its reliability. In other words, what computations can allow organisms to allocate their limited neural resources in an adaptive manner based on experience of reward and punishment? Here, we developed a general modeling framework to study how organisms endogenously allocate their attention with limited resources such that reward consumption of the organism is maximized when it interacts with the environment under uncertainty.

Formal tests of rational inattention have been predominantly tested in humans, and it remains unknown whether such influential concepts can be tested and also hold in lower-order species such as mice. Should this be the case, the availability of genetic tools (44) and large-scale cellular imaging in mice would enable the concepts studied in this work to be used to address various open questions about limited information-processing capacity (45–47) in large-scale experimental settings. This would accelerate the translation from neurobiological mechanisms to formal concepts of rational inattentive behavior in medical settings and applied economics.

To test the neurobehavioral underpinnings of rational inattention, we studied whether mice could be trained to perform a choice task requiring them to make decisions based on ordinal comparisons of orientation stimuli spanning the whole sensory space. This task decouples decisions from sensory information, rendering only measures of relative information relevant. This is essential to consider given that in ecologically valid settings, organisms typically encounter situations in which abstract choices are invariant to specific visual stimuli; for instance, when choosing between stimulus A=1 and B=2 drops of juice, the mouse must choose B, but between B=2 and C=4, the mouse must choose C and not B. Additionally, we introduced trial-to-trial variability in the reliability of sensory stimuli, and controlled the prior distribution of the sensory inputs such that it matched innate and possibly evolutionary preserved neural sensory codes in mice. During its experimental lifetime, each animal experienced novel stimulus-rewards association rules in different environments while the physical statistics of the environment and task structure remained identical. Controlling and manipulating all these components in the decision task provides a unique opportunity to study whether and how mice learn to adapt their neural coding schemes to maximize reward outcome in each environment without any cues other than trial-by-trial experience.

We found that mice indeed behave as rational-inattentive agents, by taking into consideration their information-processing limitations to develop sensory encoding strategies that lead them to maximize reward consumption. This suggests that many aspects of variable and apparently irrational behavior both in the perceptual and economic domain in fact reflect the efficient use of a limited metabolic budget to operate and interact with the environment.

## Results

### Behavioral task and behavior

We trained mice (n=7) in a two-alternative forced choice task (2AFC), in which they were presented two gratings *θ_l_* and *θ_r_* (presented on the left and right side of the screen, respectively) and had to choose the one that was more vertically oriented by turning the wheel under their forepaws and thereby moving the grating to the center of the screen (48) (Figure 1B). The two alternatives *θ_l_* and *θ_r_* are independently drawn from a prior distribution *π*(*θ*) (Figure 1A,B, see below). In order to study the role of sensory uncertainty, we randomly varied the contrast levels for each of the two stimuli across different levels on a trial-to-trial basis (Figure 1B, see methods for details). In the following, we first describe how we derived the prior *π*(*θ*) used in our study, then proceed to explain the experimental paradigm of the decision-making task, and present the descriptive behavioral results.

**Fig. 1.**
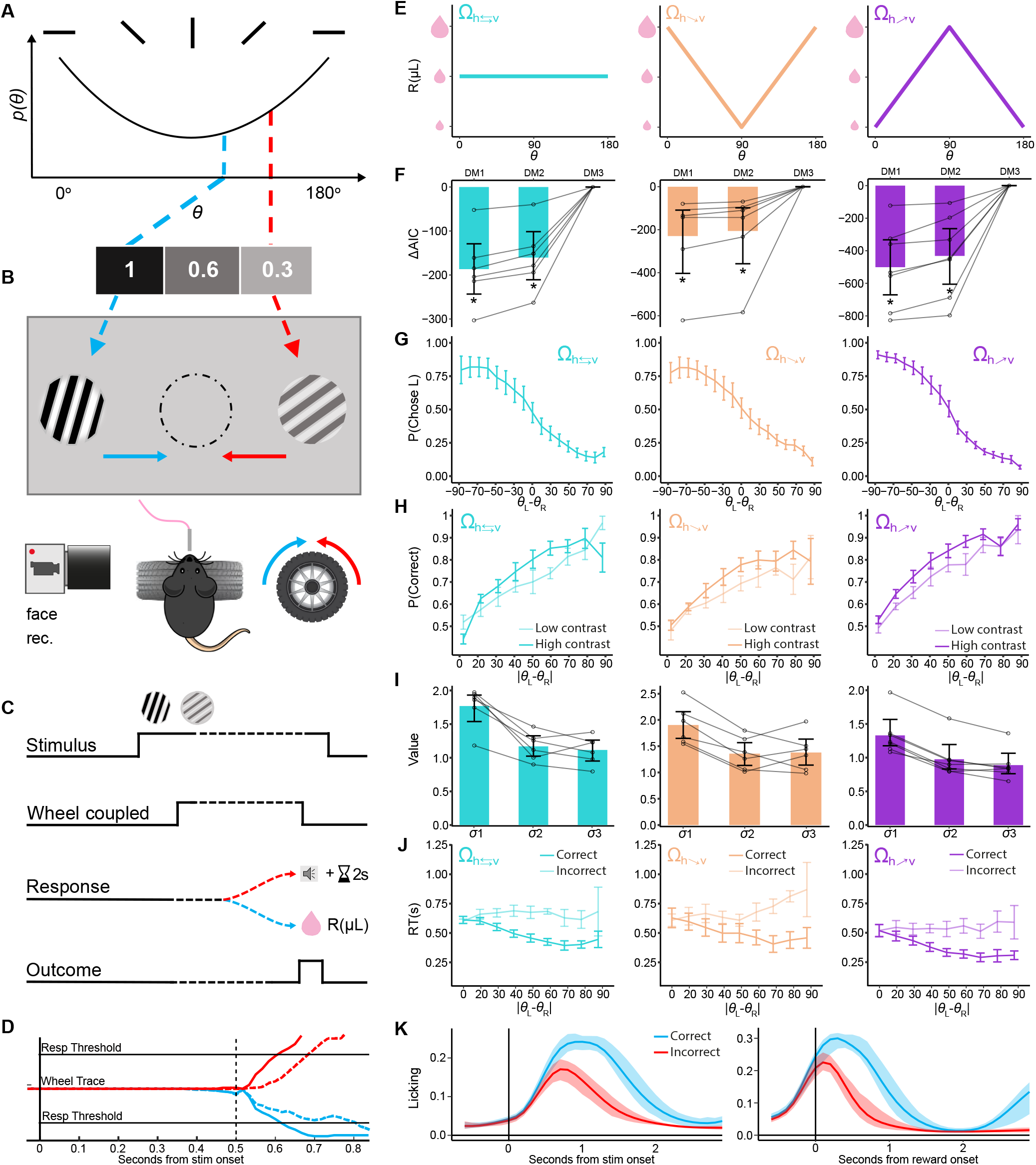
(A, B, and C) The illustration of the animal setup and procedure. (A) Two orientations were picked from a prior distribution and their contrast levels were modified independently across 3 levels. (B)The animal must turn the wheel to move the grating that is more vertically oriented to the middle of the screen to obtain a milkshake reward. (C) Trial timeline, and outcomes for correct and incorrect responses. For 0.5 seconds after stimulus onset the wheel is not coupled to the stimulus positions. (D) The median wheel trace for correct (full line) and incorrect (dashed line) trials. Trials picking right side plotted in red, and left side in blue. (E) The three stimulus-reward mapping conditions Ω considered in this study. (F) Comparison of AICs (Akaike Information Criterion) the three descriptive models (DMs) and the 3 reward mappings (Methdos). AICs from each model were subtracted from the best fitting model (DM3). Individual mice are shown with transparent gray lines. Error bars represent bootstrapped 95%-CI of the mean across mice. (*) represents models for which the bootstrapped 95%-CI AIC difference is below zero. (G) Psychometric curves for three reward mapping conditions. Mouse data is in the colour of the condition. (H) Percentage correct across difficulties for low contrast trials (semitransparent, sum contrasts < 1) and high contrast trials (solid, sum contrasts > 1). Only trials with equal contrast for left and right side stimuli are included. (I) Contrast noise assigned to each contrast level in DM3 (*σ*1, *σ*2 and *σ*3 for contrast values 0.3, 0.6 and 1, respectively). Higher weight means greater noise in the orientation estimate according to DM3. Individual mouse weights plotted as semitransparent lines. (J) Mean reaction times across difficulties for correct (solid line) vs incorrect trials (semitransparent line). In panels G-J error bars represent s.e.m. (K) Extracted licking data aligned to stimulus onset (left) and reward onset (right). Correct trials are plotted in blue, and incorrect trials in red (shaded area represent s.e.m.).

### Prior distribution approximation of stimulus orientation in mouse V1

In order to specify the prior distribution *π*(*θ*) used in our task, we investigated whether it would be possible to estimate the innate information coding of orientations in the mouse primary visual cortex (V1). To this end, we relied on a large-scale cellular imaging dataset based on simultaneous recordings from ~20,000 neurons in mouse V1, while static gratings at a random orientation were presented on each trial (47). Here, we paid particular attention to two pieces of information: (i) the distribution of preferred orientations across V1 cells, and (ii) the neural decoding orientation error of a linear decoder. Regarding the distribution of orientations, the data shows that in mouse V1, a larger proportion of neurons are dedicated to code horizontal relative to vertical orientations for static stimuli (Figure S1A). Based on this result, theories of efficient neural coding predict: (i) decoding error should be larger for orientations with less neural resources, thus higher decoding error for vertical orientations in mice, and (ii) this decoding error shape across the sensory space should be amplified for shorter exposure to sensory stimulation (19, 49). In line with these predictions, the data shows that decoding error was higher for vertical than horizontal orientations, and this pattern was further amplified in the condition with shorter stimulus presentation (Figure S1B). These results confirm that mice appear to dedicate more neural resources to encode horizontal relative to vertical orientations. We speculate that this prior distribution could have been induced in mice by evolution (or perhaps life-experience) due to their anatomy and morphology. However, this notion would need to be more formally tested in future studies, for instance as previously studied in cats, a species that appears to be more exposed to horizontal than vertical orientations in their natural environment (Figure S1C).

Based on these analyses, here we specified the distribution of orientations *π*(*θ*) such that horizontal angles were more common than vertical angles (Figure 1A). This allowed us to focus our analyses on studying how mice adopt reward-maximizing coding strategies based on specific reward-stimulus contingencies, while preserving prior neural codes that appear to be innately present in mouse early visual areas.

### Mice can perform ordinal comparisons under uncertainty in a continuous sensory space

In the 2AFC task, orientations of the two gratings were drawn from distribution *π*(*θ*) (Figure 1A). For all conditions tested in this study, mice had to choose the grating that was more vertically oriented, and received a reward *R* (a given amount of milkshake) if and only if a correct decision was made. Rewards were based on three different stimulus-reward association environments Ω (Figure 1E, Table S1). After mice reached stable behavior in a baseline decision task, mice performed between 20-50 sessions in each environment Ω (methods).

In environment Ω_*R*:*h*⇄*υ*_, mice received a fixed amount of milkshake for correct decisions irrespective of orientation (Figure 1E, left panel). This condition corresponds to the reward contingencies classically implemented in perceptual decision-making tasks. In environment 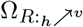, more vertical orientations yielded higher rewards for correct decisions via a linear relationship (Figure 1E, right panel). Given that vertical orientations occur less often than horizontal orientations, this context emulates ecologically-valid settings of economic decisions, where stimuli with higher value occur less often in the environment. Finally, in environment 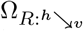 more horizontal orientations yielded a higher reward (Figure 1E, middle). Recall that in all cases the decision rule is to choose the orientation that is more vertical. Thus counter-intuitively, in any given trial in environment 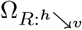, mice should choose the orientation that is mapped to the smallest reward, because the other option is less vertical and therefore leads to no reward at all.

First, we investigated based on simple descriptive models whether mice were able to follow the basic decision rule in each environment Ω (all models incorporated lapse rates and trial history effects, methods). Using a simple descriptive model (DM1) incorporating the angle difference Δ*θ* between the input stimuli in a given trial, we found that for all environments Ω, mice learned to choose the more vertical orientation (slope of psychometric curve, logistic mixed effects *P* < 0.001 for all Ω, Figure 1G). This indicates that, during their decisions, the abstract choice rule was decoupled from specific sensory information, and relative information was the relevant variable. These results are in principle in accordance with a recent study also showing that mice can apply this sort of abstract decision rules (50).

In a second model (DM2), we investigated whether stimulus contrast impacted mice decisions. As expected, we found that the lower the contrast, the noisier the decisions (mixed effects model, effect of contrast on decision noise *P* < 0.001 for all Ω, Figure 1I). In a third model, we investigated whether in addition to the influence of stimulus contrast on decision noise, the difference in stimulus contrast between the two orientations impacted mice decisions (DM3). In addition to the impact of angle difference and contrast on decision-making, we found that mice had a preference to choose the orientation with higher contrast stimulus (mixed effects model, *P* < 0.001 for all Ω). This model explained the data better than the simpler models for all mice and all environments (bootstrap 95% confidence intervals (95%-CI) of *AIC*_DM1_ − *AIC*_DM3_ < 0 and *AIC*_DM2_ − *AIC*_DM3_ < 0 for all Ω, Figure 1F). While contrast should not play a relevant role in mice decisions, below we will show that an optimal observer model can account for this apparent risk-averse behavior. Finally, we found that lapse rates were relatively low (*λ* = 0.091, 95%-CI[0.044,0.14]), suggesting that mice were engaged and followed the decision rules of our task.

For completeness we investigated whether reaction times (RTs) show the behavioral patterns often observed in both perceptual and value-based decision tasks as function of absolute evidence and trial correctness. We found that mice RTs were faster for correct responses (Figure 1J). Also the higher the absolute evidence (Δ*θ*), the faster the response for correct decisions (Mixed Linear Model, *P* < 0.001 for all Ω). These results are in line with chronometric behaviour reported across species both in perceptual and value-based decisions (51–53).

Taken together, the behavior exhibited by the mice in our study indicates that they can learn to perform decision tasks that decouple decisions from sensory information, where measures of an abstract decision rule based on relative information are relevant. Moreover, their psychometric and chronometric behavior strongly suggests that mice employed similar neural mechanisms for guiding their decisions. Next, our goal is to investigate what neuro-computational mechanisms might guide behavior under cognitive limitations.

### Efficient perception and rational inattention

We developed a framework, based on principles of optimal statistical inference, that allowed us to study how sensory systems should allocate coding resources under the assumption that there is a limit in information processing at two stages of the decision process: (i) limited precision during sensory encoding, and (ii) limited precision in down-stream decision circuits. Crucially, given that the physical prior *π*(*θ*) remains constant, these predictions may be specific to the stimulus-reward associations of a given context or environment Ω. Here it is assumed at all times that the goal of the organism is to maximize reward consumption (Figure 2A, see methods for a detailed specification of the model).

**Fig. 2.**
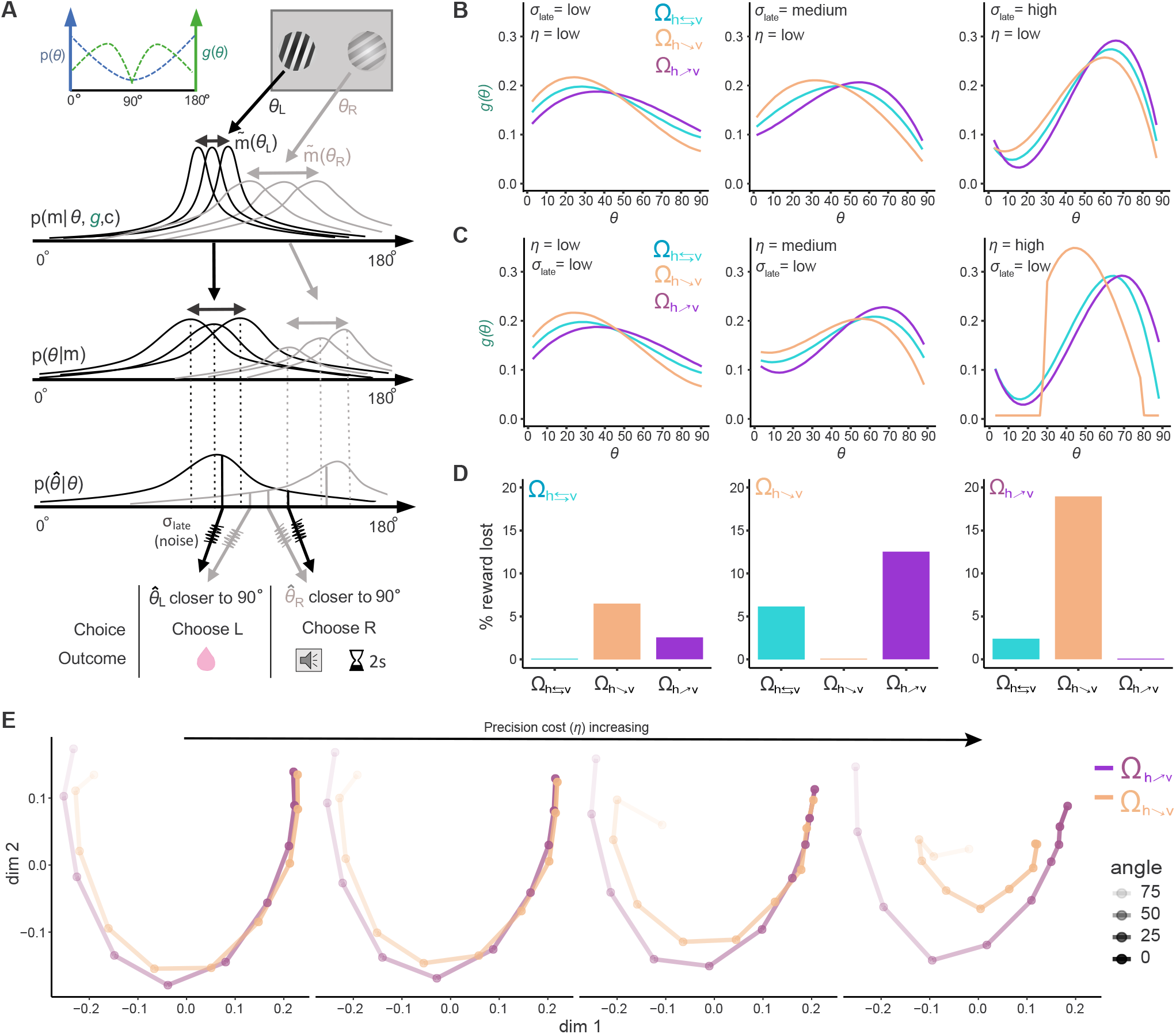
(A) Top: Prior distribution of orientations (blue) and illustration of an example gain function *g* (green). Middle: Reduced illustration of the inference model. A noisy measurement *m* is obtained for each presented stimulus *θ* according to the Likelihood function *p*(*m* | *θ*, ·). Then prior and likelihood function information are combined to derive a posterior distribution. Subsequently, the observer computes a posterior estimate 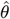 (that generates a posterior of estimates 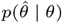 after many observations), which can be corrupted by late noise *σ*_late_. Finally the observer chooses the more vertical of the two posterior distributions. Bottom: Table of choices/outcomes dependent on noisy decoding. (B,C) The gain functions *g* which guarantee maximal reward consumption depend on the levels of both encoding cost *η* and late noise *σ*_late_. (B) The optimal solutions for *g* at three reward mappings at different noise levels. From left to right: *σ*_late_ = 0; *σ*_late_ = 0.4; *σ*_late_ = 1. Precision cost is held constant at relativel low value *η* = 0.001. Each colour represents a reward mapping condition. (C) Optimal solutions for *g* at three reward mappings at different noise levels.. From left to right: *η* = 0.001; *η* = 0.137; *η* = 0.5. *σ*_late_ = 0 at all levels. In B and C, the part of the gain function shown is from 0 to 90 degrees, with the other half, from 90 to 180, being a mirror image (as shown in A). (D) Failing to adopt efficient encoding gains *g* at for each environment Ω can lead to a substantial reward loss. Percent reward loss when parameters in optimization function for each environment Ω are swapped for another ones. Loss is plotted at *σ*_late_ = 1. (E) Geometric analysis of psychometric performance based on multidimensional scaling (MDS). Analysis based on MDS has the property that disciminibability between input alternatives in intuitively interpretable. That is, the geometric distance between a pair of situmli (each stimulus represented by a point) represents the degree of discriminability between a pair of inputs. For clarity, in this panel we present the MDS for environments 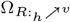 and 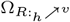 at different levels of sensory precision cost *η*. This analysis reveals: (i) discriminability at horizontal angles is higher for 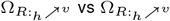, and vice-versa for vertical angles, (ii) increasing precision cost *η* dicreases discriminability in general, and (iii) for a fixed *η* level, discriminability is higher for environment 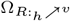 relative to 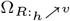.

Given that noisy communication channels always lose information during transmission, we argue that it is more efficient for the brain to adapt to the reward-maximizing rules of a particular environment at the earliest stages of sensory processing. This notion is supported by evidence suggesting that early sensory systems represent not only information about physical sensory inputs, but also non-sensory information according to requirements of a specific task and the behavioral relevance of the stimuli (54–56). Thus, a key assumption of our framework is that, despite the fact that the physical prior distribution was held constant for all environments Ω, sensory codes adapt to environmental demands, possibly via feedback schemes (57, 58).

In brief, we assume that in every trial a mouse obtains a noisy measurement *m* independently for each input orientation *θ*. We model the noisy measurements using the von Mises distribution

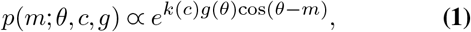

where the precision of the measurement is determined by two multiplicative factors: (i) *k*(*c*), which is function of the contrast level for a given orientation, and (ii) a normalized limited-resource function *g*(*θ*), determining how many resources the system allocates to a particular segment of the orientation space (methods).

On each trial, the agent computes a posterior distribution *p*(*θ* | *m*) by combining the physical environmental prior distribution *π*(*θ*) with the likelihood of the measurement *p*(*m* | *θ*) via Bayes rule. Then the agent applies a decoding rule in order to obtain a posterior estimate 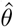, which we assume to be the expected value of the posterior distribution *E*[*p*(*θ* | *m*)], also known as Bayesian least squares (BLS) estimator. We assume that on each trial, the mouse independently estimates 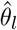 and 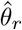 for the input orientations *θ_l_* and *θ_r_*, respectively, and then makes a decision based on the abstract decision rule in our task: choose the more vertical angle.

If we constrain *g*(*θ*) > 0 for all *θ* and assume a cost *K* generated by the use of metabolic resources to encode visual information (35), one can formulate a well-constrained optimization problem in order to find the optimal allocation of resources in the orientation space for: (i) a given physical environmental prior *π*(*θ*), (ii) a given contrast response function *k*(*c*), (iii) reward outcomes associated to decision-outcome rules in a given context or environment Ω, and (iv) downstream noise *σ*_late_ that is not related to sensory encoding. Formally, the goal is to find a resource allocation *g*\*, and the maximum allowed precision/neural-activity *k*_max_ (with an underlying contrast response function *k*(*c*), see methods) such that

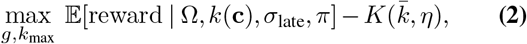

where **c** is the set of contrasts that the animal experiences. We model the metabolic cost *K* as a function of the average precision 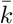 that the agent invests on solving the inference problem, where *η* > 0 indicates how much the metabolic cost scales with average precision 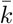 (methods). We argue that this choice of cost function is reasonable when applied to the visual system, given that in our model sensory encoding precision (Eq. 1) can be directly related to the amount of neural activity and its associated variability (13) (methods, Figures S2 and S3). Moreover, it has been recently shown that the relationship between neural activity and energy expenditure in the brain is nearly linear (59).

Here it is important to highlight that an important feature of the rational inattention framework is that allocation of sensory precision (*g*(*θ*) and *k*_max_) does not need to be assumed or manually fitted—as it is often the case in plain Bayesian frameworks were likelihood functions are assumed or manually fitted—but emerge endogenously by assuming an average precision cost *η* and downstream precision *σ*_late_, along-side the associated (reward) loss that the organism experiences (Methods).

### Optimal resource allocation depends on both sensory encoding precision and down-stream noise

We studied whether the optimal limited-resource allocation function *g* depended on the amount sensory precision cost *η*. Additionally, we also studied the dependence of *g* on sources of down-stream noise *σ*_late_, as recent evidence suggests that limits of sensory perception in mice might be also related to downstream neurocomputational imprecision (47), possibly related to computational limitations of downstream decoders and other forms of irreducible noise in the system (33). Here, we would like to emphasize that a key feature of our study is that for all three different stimulus-reward environments Ω, the physical prior *π*(*θ*) was always the same, and therefore, potential differences in resource allocation *g*(*θ*) must be related to internally improving sensory representations of the learned associations that maximize reward expectation. In order to derive the optimal *g* functions for a particular combination of *η* and *σ*_late_, we used the exact distribution of stimuli inputs used in the mice experiments.

We found that in general for low levels of downstream noise *σ*_late_ and relatively low costs in encoding precision *η*, more resources should be allocated to segments of the orientation space with higher prior *π*(*θ*) density (Figure 2B, left panel). This result is in line with previous studies suggesting that in the limit of low sensory noise (i.e., *η* → 0), encoding precision should be higher for segments of the orientation space with more density (19, 60). For instance, it can be shown that for the case of perceptual tasks with constant reward delivery per correct trial (i.e., Ω_*R*:*h*⇄*υ*_ environment in our study), disrimination thresholds are inversely proportional to the prior distribution, but crucially in the low-noise regime (19, 49). However and surprisingly, as *σ*_late_ increases the predictions of these analytical solutions breakdown, and agents should start to be myopic to segments of the sensory space where the prior’s density is high (Figure 2B, middle and right panels). Similarly, assuming no downstream noise (i.e., *σ*_late_ = 0), but instead varying the levels of sensory noise reveals a similar but qualitatively different pattern of optimal resource allocation strategies as a function of sensory precision costs (Figure 2C, Figure S2,3,4). Thus, these predictions reveal two important features of the rational inattentive agent. First, optimal resource allocation does not only depend on the level of encoding precision of sensory signals, but also on stimulus independent downstream noise. Second, long-held conceptions that neural resources should always be allocated to spaces with higher prior density do not necessarily hold beyond the low-noise regime.

Why are more resources allocated away from high prior density spaces when precision cost and downstream noise are relatively high? On the one hand for the case of high precision cost, low levels of encoding noise generate higher levels of Bayesian attractive biases. Thus, optimal solutions of the rational inattention model reveal that if the system has the opportunity to flexibly modulate its sensory gain, then it pays off to put more weight at places of the sensory space that avoid poor discriminability due to overall attractive biases. In order to provide a better intuition behind this result, we carried out geometric analyses of psychophysical performance based on multidimensional scaling (MDS) (Figure 2E and Figure S5). MDS reveals that the effect of attractive biases for high precision costs are compensated by expanding the discriminability of orientations located at intermediate levels of verticality. On the other hand, for the case of high down-stream noise, augmenting precision at the edges of this space does not increase the discriminability of neighbouring orientations beyond the most vertical and most horizontal orientation. That is why for relatively high levels of noise there is preference to allocate more attentional resources to rather intermediate levels of cardinality (Figure 2E and Figure S5). While this pattern is similar for all environments Ω considered here for a given combination of *η* and *σ*_late_ (Figures 2B,C and Figure S2), there are important differences in the resource allocation functions *g* between environments. We found that resource allocation has a tendency to be relatively higher for more vertical angles in environment 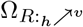 (Figure 1E, middle panel), however this requires sacrificing precision at more horizontal orientations. This pattern reverses for environment 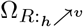. This result intuitively makes sense, given that in environment 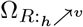, more vertical angles deliver more reward, and therefore mice are better off at giving up encoding precision for horizontal orientations even if they occur more often. Another key prediction of the rational inattention framework is that for environment 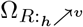 the system should allocate more resources to solve decision task relative to 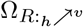 (this is evident in the MDS analyses where nodes of the manifold are more separated in 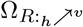 relative to 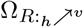). This prediction emerges in the rational inattention framework because the relative allocation of resources under our inference framework leads to slightly higher reward expectation in environment 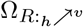 relative to 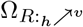. We also considered rational inattentive model that does not allow a variable and adaptive gain function *g* across the sensory space, but instead we assume it to be constant. While this model captures the effect where in environment 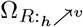 the system allocates more resources to solve decision task relative to 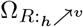, this model does not account for tradeoffs in discriminability between horizontal and vertical spaces across environments (Figure S6). Moreover, the MDS manifold is more spread for the *g*-variable model in relation to the *g*-constant model, thus suggesting higher efficiency in the *g*-variable (compare manifold spread in Figures S5 and S6).

Finally, we investigated what is the actual benefit of applying the most efficient resource allocation in a given environment. Using average parameters of model fits to the mice data (see below), we found that agents could lose around 10-20% of reward, if resources are not optimally allocated (Figure 2D). This confirms that efficient resource allocation under uncertainty and cognitive limitations is critical to maximize reward.

### Adaptive rational inattention in mice

First, we investigated whether the rational inattention model that endogenizes *g*(*θ*) (*g*-endog model) provides a better account of the mice data than the best descriptive model (DM3, Figure 1F). We found that the *g*-endog model provides better fits to the data for all environments (bootstrap 95%-CI of *AIC*_RI_ − *AIC*_DM3_ < 0 for all Ω, Figure 3A). Notably the rational inattention model has the same number of parameters as the best descriptive model. Therefore, the quantitative differences in goodness of fit between the two models is not related to model complexity. Inspection of the *g* functions across mice for each environment follow the counter-intuitive qualitative predictions of the theory for non-negligible sensory and late noise in our task: More resources are allocated to angles close vertical relative to horizontal orientations (Figure 3C). Additionally, more resources are allocated to vertical orientations in environment 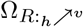 (higher reward for more vertical angles, Figure 1E) relative to the other environments, while the opposite is the case for 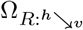 environment (cluster-corrected P<0.05; Figure 3C). These results strongly suggest that the manner in which mice allocate attention in light of their limited coding resources—in the presence of encoding and precision noise—follows the signatures of the resource-limited observer model. Additionally, we found that the levels of apparent risk aversion behavior observed in the descriptive model (DM3, see above) can be largely explained by the rational inattention model (Figure 3B).

**Fig. 3.**
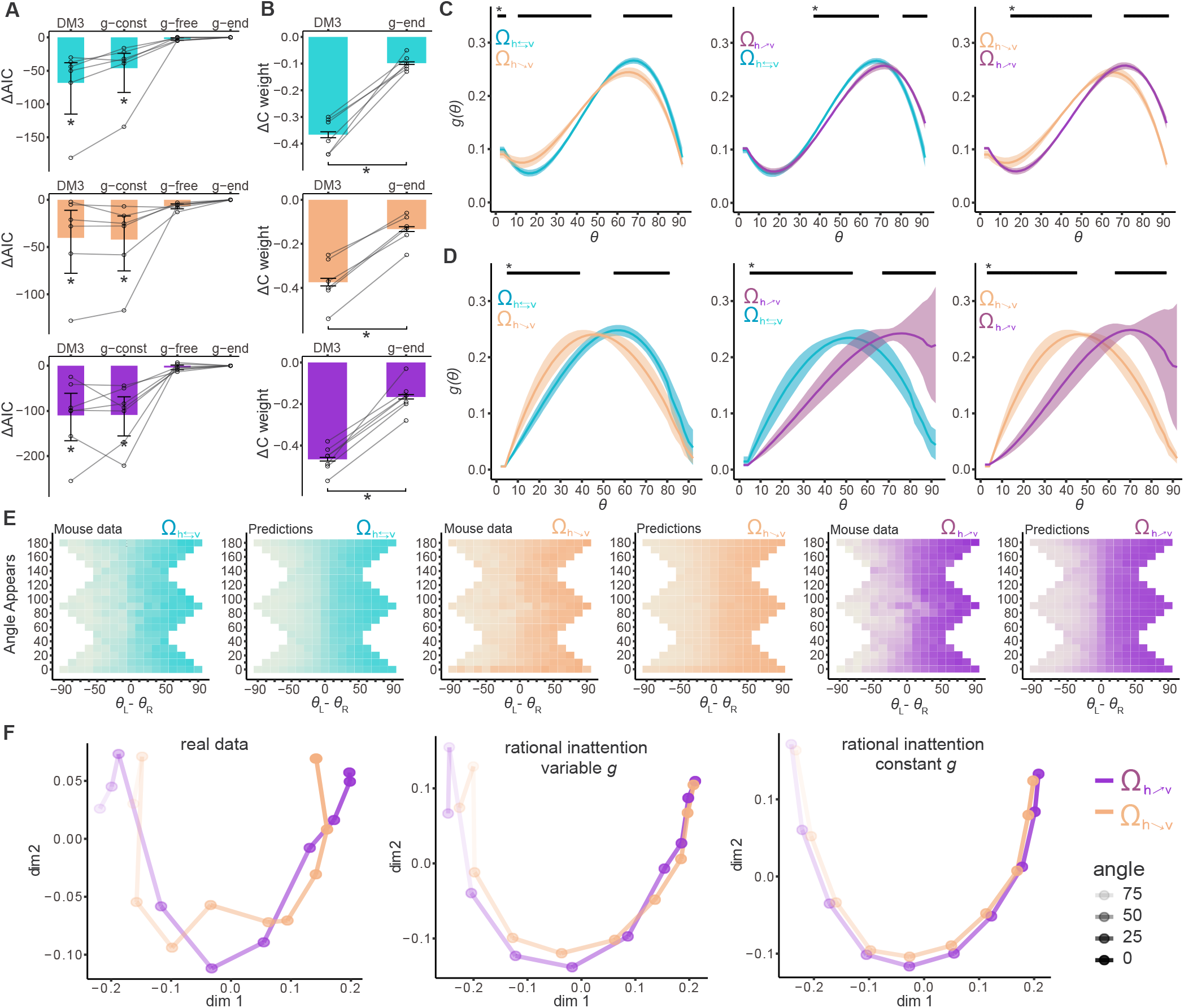
(A) Model comparison between based on the AIC reveals that the rational inattention model provides the best account of the behavioral data for all enviroments Ω. The figure shows the difference in AIC relative to the *g*-endogenized rational inattention model (*g*-end). Error bars represent the bootstrap 95%-CI across mice, and individual mice are shown with transparent lines. (*) Denotes bootstrap 95%-CI significant difference vs the g-endogenous model. There is no difference between the model that endogenizes *g* and the non-endoginized *g* model, suggesting that model complexity that arises by freely fitting *g* is not necessary. (B) Probit wights capturing the influence of contrast difference between the two stimuli on choice. Large part of the apparent risk aversion observed in the best descriptive model DM3 (i.e;, preference to choose the stimulus with higher contrast) is captured by the rational inattention model. Error bars represent s.e.m., and individual mice marked by transparent lines. Asterisk denotes significant differences (mixed effects model, P<0.001). (C,D) Pairwise comparisons of resource allocation functions *g*(*θ*) for each reward environment. Black lines on top (marked with an asterisk) signal significant differences as determined by bootstrap 95%-CIs of within mouse differences. Panel C shows results for *g*-endogenised model and panel D for the non-endogenized model. See Figure S8 for individual mice data. (E) Heatplots for each reward environment showing probability of choosing the left stimulus at each possible orientation appearing in the study vs. the angle difference (trial difficulty). Model predictions and real mouse data are shown. Data and model predictions show overall good agreement. See Figure S7 for data split based on different contrast levels. (F) MDS analyses of real data and model fits reveal that the rational inattention model allowing adaptive *g* across the sensory space has a psychophysical geometry that closely resembles the real data. On the other hand, the rational inattention model that assumes a constant gain *g* cannot account for various discriminability features observed in the data.

In order to make sure that the shapes and differences in resource allocation *g* are not a product of the endogenous solutions of the optimal models, we fitted a version of our inference model that does not endogenously restrict the shape of *g*, but is fit in a model-free fashion (methods). This model reveals two striking results: First, the qualitative shape of resource allocation *g* across environments is similar to the optimal solutions (more resources are allocated away from segments of the sensory space with high prior *π*(*θ*) density). Second, mice allocated their limited coding resources in a way that resembled the signatures of reward-maximization in each environment Ω. More specifically, we found that (i) in environment 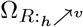 (more reward for more vertical angles) resource allocation was relatively higher for vertical angles, (ii) this pattern reversed for 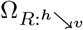, and (iii) resource allocation in environment Ω_*R*:*h*⇄*υ*_ was located in between the other environments (see pairwise comparisons in Figure 3D, P<0.05 cluster-corrected). Crucially, model comparison between the *g*-endogenous model and the *g*-free model reveals that the complexity of freely capturing the shape of resource allocation *g* is not necessary (bootstrap 95%-CI of *AIC*_g-endog_ − *AIC*_g-free_ < 0 for all Ω, Figure 3A). In addition to these quantitative results, our model also accounted for various qualitative features of mice behavior (Figure 3E, and Figure S7) as it is also evident based on MDS analyses, which reveal that the *g*-endog model closely resembles the geometry of psychophysical performance of the real data (Figure 3F). Here, two key predictions of the rational inattention model are observed in the behavioral data: (i) geometric distance of the nodes in the MDS manifold at horizontal orientations is shorter for environment 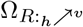 relative to 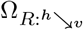, and (ii) geometric distance is larger for environment 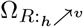 relative to 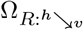 (see Figure 2E). These results suggest that mice indeed adopt efficient rational-inattentive perception according to the reward contingencies of a given environment.

Finally, We compared the results of the endogenized model (*g*-endog), with an ideal observer model in which resource allocation was uniform across the whole sensory space (*g*-const model). We found that the *g*-endog model accounted better for the data than the *g*-const model for all mice and all environments Ω (bootstrap 95%-CI *AIC*_g-endog_ − *AIC*_g-const_ < 0 for all Ω, Figure 3A,F). These results strongly suggest that assumptions in ideal observer models where uniform likelihood functions are adopted (11, 61, 62)—mainly for mathematical convenience—might not always be warranted.

### Adaptive rational inattention via reinforcement learning

The rational inattention model implemented above assumes that mice have already adapted to a permanently relevant environment Ω. Hence, the optimal allocation of resources *g* remains fixed across trials in a given environment. Therefore, this model does not explain how mice may re-allocate their limited coding resources via trial-to-trial experience. In order to incorporate this fundamental aspect of behavior in our model, we implemented a class of reinforcement learning (RL) mechanism allowing dynamical updating of the resource allocation function *g* using operations that appear to be supported by biological neural systems (Figure 4A, see methods for details). In brief, we assume that the animal updates—based on experience—a distribution of rewards associated to sensory information, for instance in the VTA (63), or any other downstream circuit performing similar operations. After a mouse makes a decision, the animal updates the reward distribution using a form of distributional updating (see methods). Crucially, the strength of update (i.e., learning rate) is dynamically adjusted according to the confidence level of the decision just made (64). Additionally, based on compelling evidence suggesting that humans learn differently from positive and negative outcomes (65, 66), we investigated whether our mice would show a similar behavior in our task by allowing separate learning rates for correct and incorrect decisions. In the final step, we assume that the brain readjusts the resource-limited sensory gain *g* (possibly in early sensory areas) via a top-down divisive normalization operation (67, 68).

**Fig. 4.**
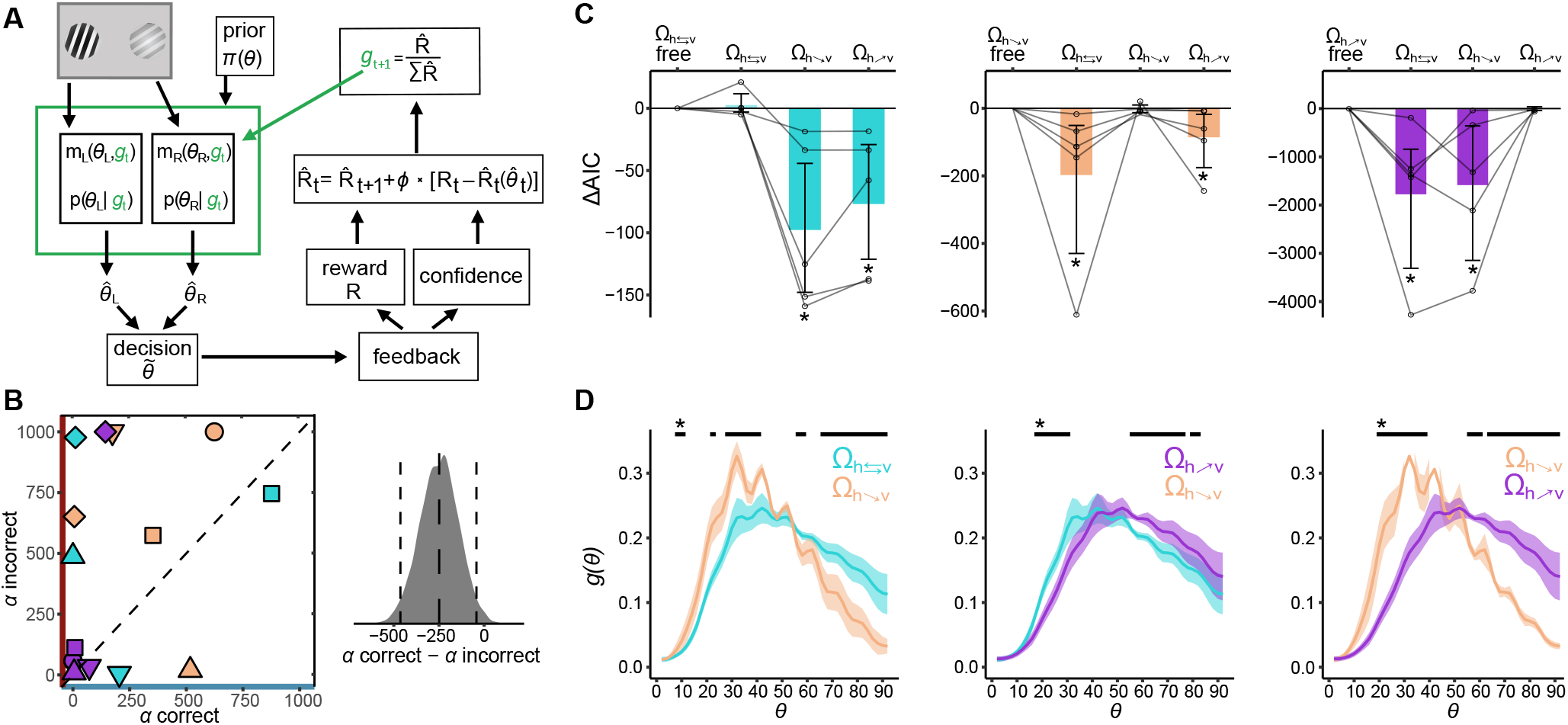
(A) Sketch of the reinforcement learning (RL) model. In brief, after a the observer chooses angle 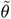, the gain *g* function is updated following a divisive normalization operation that depends on reward received *R*, the level of confidence in the decision and a distributional update of the reward distribution vector 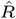. (B) Left: Learning weights *α* for correct and incorrect trials. Individual mice shown with different symbols, and color-coded for condition (see Figure 1D). Right: Distribution of bootstraped differences of *α* for correct and *α* for incorrect trials, dashed lines show the mean and bootstrapped 95%-CIs. This result suggests that learning rates are higher for correct choices vs incorrect choices (recall that our model, the learning rates *α* are divisive, and therefore, lower values indicate higher learning rates). (C) AIC difference between the *g*-endogenized model and the learning model for each environment Ω. Learning parameters are in each subplot from left to right swapped for each condition shows that there were no difference between the endogenized rational inattention model and the learning model in its corresponding environment Ω. Semitransparent lines show the AIC differences for each mouse and error bars represent the bootstrapped 95%CI across mice. Significantly different models denoted by an asterisk. (D) Resulting gain functions *g* from the RL model averaged for the last 500 trial in each environment Ω for each mouse and then averaged across mice. Error bars represent s.e.m., and black lines on top show significant differences evaluated by studying whether the bootstrapped 95%-CIs of within subject differences do not cross zero.

If it is the case that the RL model described above converges to the static rational inattention model, we expect a similar allocation of attentional resources *g* across environments for both models. We found that the average resource allocation functions *g* over time showed nearly identical patterns of those found in the static ideal observer model (Figure 3D). Moreover, we found that the set of RL parameters that were fit to each condition improved model fits specifically for the corresponding environment (Figure 3C). This suggests that parameters controlling the dynamics of reward-sensory learning are specific to the reward-maximizing rules of a given context. Additionally, the difference between the static rational inattention and RL models was not distinguishable across mice for all environments Ω (Δ*AIC*(Ω_*R*:*h*⇄*υ*_) = 2.8[−3.4, 12.4]; 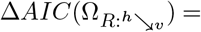 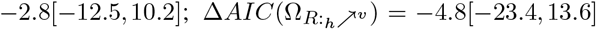, Figure 3C). Given that the rational inattention model fits the data better than all other models tests here so far, these results suggest that the RL model approximates closely to the solution of the optimal model (bootstrap 95%-CI of Δ*AIC* < 0 between RL and all other models except the rational inattention model).

A recent intriguing modeling study found that when updating the value of a chosen options, estimates must be revised more strongly following positive than negative reward prediction errors, as this guarantees that reward expectation is maximized (69). Crucially this effect is amplified when decisions are corrupted by late noise. Given our finding that downstream noise has an important influence in the allocation of limited resources and consequently on decision behavior, we hypothesized that similar to humans (65), learning rates in mice are also higher for correct relative to incorrect decisions in our adaptive resource allocation model. In line with these predictions, we found that learning rates were generally higher for correct relative to incorrect decisions (Δ*α* = −249.0, bootstrap 95%-CI[−465.6, −41.7], Figure 3B), thus suggesting that apparent irrational learning policies might indeed lead to reward-maximizing strategies.

### Rationally-inattentive neural population codes

We investigated allocation of neural resources of our sensory task in a model that is more closely related to a possible biological implementation. To this end, we implemented a neural network of V1 neurons with Poisson spiking statistics. Thus, an advantage of studying such an implementational architecture is that we can directly study the trade-off between reward intake and the metabolic costs associated to the generation of action potentials in the network (70). Here we provide a brief description of the model alongside the main results (Figure 5A, see methods for more details).

**Fig. 5.**
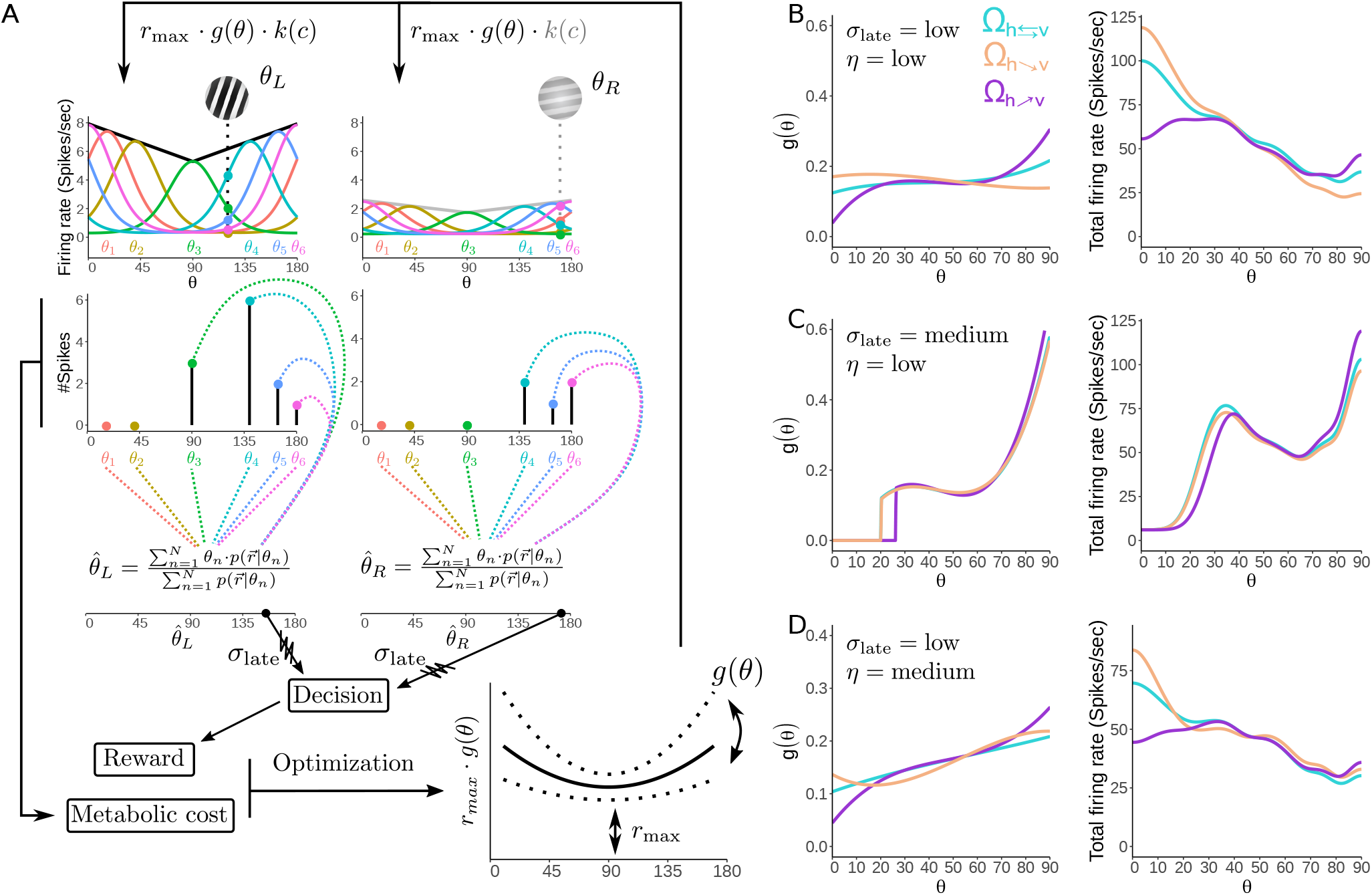
(A)Schematic of the neural network inference process. Top row: the preferred stimuli *θ*_n_ of *N* = 30 neurons are spread through orientation space proportional to the prior distribution (for visualization purposes only 6 neurons are depicted). The neurons follow a bell-shaped activation pattern which is modulated by the maximum firing rate *r*_max_ and the resource gain function *g*(*θ*). Neural activities are in turn modulated by a contrast response function *k*(*c*). The two input angles *θ_L_* and *θ*_R_ activate independent and retinotopically specific neural networks. Note that in this example the low activation of the neurons for *θ_R_* is a result of low contrast level. 2nd row: the activation of Poisson neurons leads to a number of spikes for each neuron. 3rd row: the BLS estimates of the input angles 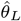 and 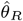 are computed by summing over the preferred orientation of the neurons multiplied by the likelihood. Bottom row: a noisy decision is made based on the BLS estimates of the input angles. The decision leads to a reward conform the choice rule and the reward environment. Both the reward and the metabolic cost originating from the spikes feed into the optimization process which determines the shape of *g*(*θ*) and the value of *r*_max_. (B,C & D) Optimal solutions of the limited resource function *g*(*θ*) for different values of the metabolic cost of a spike and different levels of late noise *σ*_late_ for the different environments Ω (left) and the resulting total activation (as a function of input angle *θ*) (right). (B) Low external noise and low metabolic cost. (C) Medium external noise and low metabolic cost. (D) Low external noise and medium metabolic cost. Note that the scale of the y-axis has changed in D) compared to B) and C).

Similar to the algorithmic model, the firing rate of the neurons is determined by two multiplicative factors: (i) maximum activity *r*_max_, determining the maximum firing rate of the neurons (which is also modulated by contrast input), and (ii) *g*(*θ*) which determines the firing-rate gain of the neurons.

Based on this architecture, it is possible to show that the BLS estimate of the input *θ* can be computed by a sum over the likelihood functions based on the neural activity generated across the network 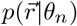 weighted by the preferred stimuli of the tuning curves *θ_n_*. Given that the physical prior distribution remained constant across the different environments Ω, we assume that the neural network incorporates this knowledge in its architecture, such that biologically-plausible computations can be implemented during encoding and decoding operations of the network. One way to achieve this is by assuming that the locations of peak sensitivity of the neurons are spread through orientation space proportional to the prior (60, 71). Conveniently, this strategy can be incorporated by the BLS estimator via the spread of the preferred stimuli locations of the neurons. We assume that these computations occur in parallel by retinotopic specific networks that independently compute the BLS estimates of the two input angles 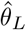 and 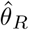. As in the algorithmic model, we allow the possibility to corrupt the estimators with down-stream noise (*σ*_late_). The decision is associated to a reward outcome conform the decision rule and the reward contingencies of the environment Ω. Crucially, we can directly relate the metabolic cost expected amount of spikes generated by the network, which we can assume to be proportional to some metabolic cost per spike *η* (72). As in the algorithmic model, the goal is to find a resource allocation *g*\* and a maximum allowed activity *r*_max_* such that the trade-off between reward and costs is optimized (see methods and Figure 5A).

Mirroring the results of the algorithmic model, we find that the allocation of resources in the network (measured as firing rates elicited for each input stimulus in the sensory space) depends on encoding noise, late noise, and the environmental context Ω (Figure 5B,C,D). For low levels of downstream noise *σ*_late_ more resources are allocated to segments of orientation space that correspond to high prior *π*(*θ*) density (Figure 5B,D). This intuitive result is not found when *σ*_late_ is increased, instead most resources are located at those regions of orientation space that correspond to low prior density (Figure 5C). Moreover, the neural network will spend less resources in the case that the cost of spiking *η* is high (i.e., average firing rate decreases, Figure 5D). Thus, this model makes testable predictions for the spiking behavior of neurons in the face of prior densities, stimulus-reward association contexts, and varying levels of encoding and late noise, that can be studied in future imaging studies.

### Arousal-linked efficient regulation of behavioral variability

Past research has identified that systems regulating arousal levels—such as the locus coeruleus–norepinephrine (LC–NE) system—have a considerable impact on behavioral variability (73, 74). For instance, it has been shown that large pupil baseline (i.e., pupil dilation before trial onset) is associated to slower and less accurate perceptual decisions (75). Moreover, in behavioral paradigms involving uncertain environments, large pupil baselines predict exploratory behavior (76, 77). These findings support the idea that tonic LC-NE modes produce an enduring and largely nonspecific increase in behavioral sensitivity, which promotes flexible and exploratory control states (77).

Based on this evidence, we hypothesized that as for the case of humans, baseline pupil dilation in our task is predictive of task disengagement, that is, more incorrect and slower responses. Based on camera tracking, we analyzed the dynamics of pupil dilation relative to trial onset and its relation to reaction times (RTs) and correct responses (Figure 6C,E, methods). In line with our hypothesis, we found that higher levels of pupil baseline were associated with more incorrect and slower responses in the upcoming trial for all environments Ω (Bootstrap 95%-CI<0 for all Ω from −0.75 to 1.5 s relative to stimulus onset, Figure 6B). Given that RTs are usually related to the degree of trial correctness (Figure 1J), we investigated whether the effect of pupil baseline on RT was also present when splitting the data in correct and incorrect trials. We found that the effect of RT was robustly present for both correct and incorrect trials (Bootstrap 95%-CI<0 for all Ω from −0.75 to 1.5 s relative to stimulus onset, Figure 6E). These results confirm that during high levels of tonic arousal, as in humans, mice show signatures of task disengagement. But, what mechanisms support this arousal related behavioral variability?

**Fig. 6.**
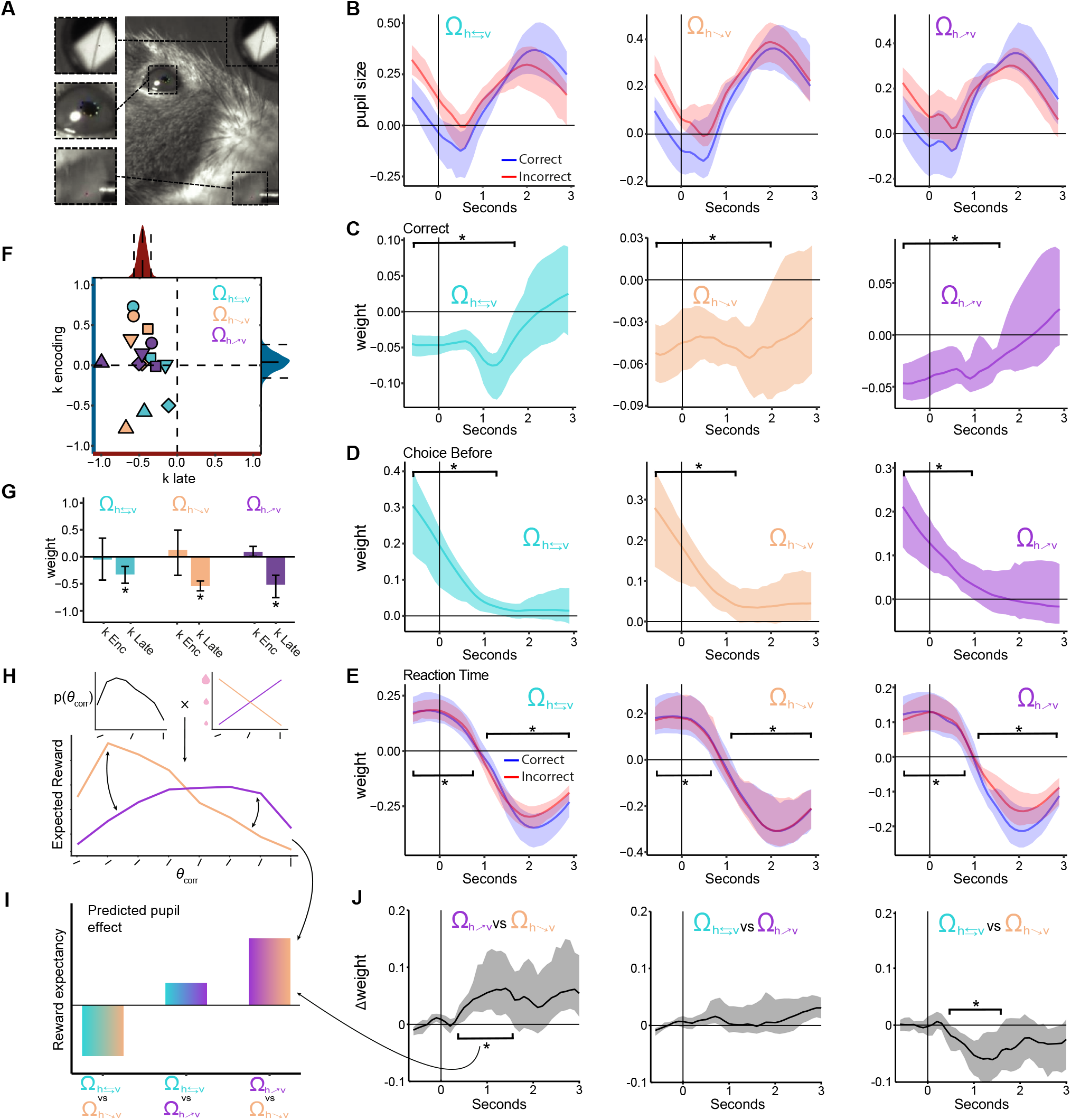
(A) Pupil dilation was extracted from videos acquired during behavioral performance using DeepLabCut. Other relevant variables can be detected such as trial onset (top window), pupil size (middle), and licks (bottom) (B) Average pupil size for each reward environment Ω. Blue line shows pupil size for correct and red line for incorrect trials. Shaded area shows 95% bootstrap CIs. (C-E) Coefficient estimates eastimated for each time point based on multiple linear regressions with pupil size as the dependent variable and multiple trial parameters as predictive variables (see Methods). Panels show the coefficient estimates over time for trial correctness, correctness on a choice before, and reaction time (where multiple regressions were ran separately on correct and incorrect trials). In panel E it can be appreciated that baseline pupil size predicted slower decisions in the upcoming trial irrespective of trial correctness. (F) Baseline pupil dilation directly impacts downtream noise precision but not sensors precision. Regression weights of sensory precision (k-encoding) vs. downstream choice precision (k-late) as a function of pupil dilation before trial onset. Individual mice shown with different symbols, and color-coded for condition (see Figure 1D). Bootstrapped distributions projected parallel to relevant axis with mean and 95% CI shown in dashed lines. (G) Bootstraped means and 95% CIs for sensory and downstream noise parameters in each reward environment Ω. (H) Sketch showing how values for Expected reward were extracted for each correct side angle. The prior distribution of correct side angles is multiplied by the reward mapping of the respective reward environment Ω. (I) Sketch of the predicted effect of reward expectancy on pupil size when comparing between environments (i.e., the interaction effect). Arrows pointing to the furthest right bar illustrate the comparison between 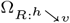 and 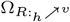 reward environments, as otherwise shown in H and J. (J) Pairwise reward environment difference in weights of expected reward to phasic pupil response over time from trial onset. All time bins marked with asterisks are significantly different from 0 (as determined by 95% bootstrap CIs). Interaction patterns closely follow the interaction predictions, suggesting that arousal systems (linked to pupil dilation) potentially encode expected reward distributions in a sensory-specific manner. pupil responses in each environment and individual mice data are shown in Figure S9.

Recent theories of the LC–NE system suggest that windows of high arousal might be related to adjustments in learning representations (e.g., in situations of high volatility where learning rates must be up-regulated (78, 79)). But, where does this trade-off occur during decision-making? An advantage of our rational inattention model is that it allows us to separate sensory encoding precision from stimulus-unspecific downstream noise. Therefore, we adapted our model to obtain a joint readout of change in sensory and late noise as a function of pupil baseline (crucially, we can show that the contribution of the two noise sources is identifiable). Strikingly, we found that for all environments Ω and all mice, higher levels of pupil baseline resulted in a consistent reduction of downstream choice precision *k*_late_ (Δ*k*_late_(Ω_*R*:*h*⇄*υ*_) = −0.33% 95%-CI:[−0.48, −0.18], 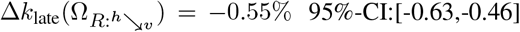, 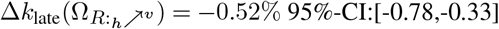; individually estimated MLE 95%-CI<0 for each mouse in each environment Ω), but not related to changes in sensory precision (Figure 5F,5G).

Given our finding that positive prediction errors are associated with higher learning rates (Figure 4B), and previous reports on the positive relationship between learning rates and tonic arousal (78), we hypothesized that following positive prediction errors, pupil baseline should be elevated. In line with this prediction (and after controlling for confounding factors), we found that positive prediction errors were related to elevated pupil baseline of the following trial (Figure 6E). Conversely, higher levels of phasic pupil response were related to faster reaction times in the current trial (Figure 6E). Thus, these findings suggest that arousal systems in the brain balance the costs associated to larger learning updates with lower precision in downstream circuits while preserving sensory fidelity. This supports previous suggestions of nonspecific increase in behavioral sensitivity during high tonic arousal, which in turn promotes flexible and exploratory control states (76, 77).

### Signatures of reward distribution encoding in arousal systems

Previous work provided evidence that the arousal system has important and computationally complex roles in rationally regulating the influence of incoming information on beliefs about dynamic and uncertain environments (78). However, it remains unclear whether the arousal system signals the expectation of rewards tied to specific sensory information (which is often distributed across the whole sensory space), irrespective of the expectation of physical sensory signals. Our experimental design allows the possibility to study these arousal-linked computational mechanisms given that the distribution of sensory signals remains identical across the different environments Ω, where the only difference across environments is the learnt stimulus-reward association spanning the sensory space.

In order to study this possibility, we investigated phasic pupil responses as a function of sensory-specific reward expectation. First, we obtained the expected reward of each correct-side stimulus by multiplying the frequency of occurrences of these stimuli with their reward value within each reward environment Ω (Figure 6H). If it is true that pupil responses signal the distribution of reward expectation in an adaptive manner, then we expect an interaction of the pupil responses linked to sensory specific reward expectation across environments (Figure 6H).

We used these values in a per-time-bin linear regression analysis on phasic pupil responses. Critically, we controlled for confounders such as trial difficulty (absolute angle difference), reaction time, correctness on a trial, the probability of appearance of an angle, and licking by including them as regressors. In a within mouse, pairwise comparison, we find a significant difference between the egression weights assigned to expected rewards in different reward environments Ω as predicted by our postulated computations (Figure 6I,J). A positive interaction between 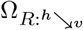 and 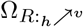 is expected, due to the inverted angle-reward mappings between them (Figure 6H,I). The phasic pupil activity reveals such interaction patterns (Figure 6J left panel, P<0.05 cluster corrected). The opposite pattern is predicted by the comparison between 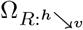 and Ω_*R*:*h*⇄*υ*_ (Figure 6I), which is also confirmed by the phasic pupil responses (Figure 6J right panel, P<0.05 cluster corrected). On the other hand, the predicted difference between 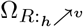 and 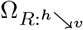 is relatively small with a slight tendency in the positive direction (Figure 6I). This prediction was also evident in our data (Figure 6J middle panel).

This set of results indicate that pupil size increases with rarity and size of the reward indicated for a give set of sensory stimuli in a given trial. This suggests an indirect link between arousal systems, such as the noradrenergic system of the LC, and reward expectancy, which might guide learning and redistribution of sensory resources via dopaminergic systems. Interestingly, a recent investigation showed that dopaminergic responses are amplified by rarity and size of reward received in a trial (80). In addition, they show that pupil dilation is also sensitive to such rewards after feedback, thus, pointing to a fundamental interaction between noradrenergic and dopaminergic systems to guide efficient learning and decision-making behavior.

## Discussion

When organisms face a decision, choices must be made based on imprecise perceptions arising from the information-processing limitations of the nervous system. We developed a framework to study how organisms allocate attention during sensory encoding, such that there is an advantage to being stochastic, and in some cases myopic to incoming sensory information, because the encoding strategy guarantees that metabolic investment in more precise awareness leads to maximal reward consumption. We formulated the framework as generally as possible while conforming closely with neurobiological mechanisms so that the nature of the neural-coding constraints need not be fine-tuned independently for each particular context. Additionally, we show that the information-processing strategies of a rationally inattentive agent are dependent on two main factors: First, the system has limited capacity to process information, not only at the early stages of sensory encoding, but also at late stages of the decision-making process. Second, strategies depend on context; in our work, context was defined by the stimulus-reward associations in a given environment, but the stimulus prior remained constant at all times. We found that mice behave as rationally inattentive agents: they take into consideration their information-processing limitations to develop efficient sensory encoding strategies that lead them to maximize reward consumption.

Within the specific structure of our decision task, these rational information-processing strategies lead to results that seem surprising and also appealingly intuitive. When information-processing resources are low for both sensory encoding and downstream decision computations, the rationally inattentive agent is relatively myopic to portions of the stimulus space where the density of the prior is highest. This result goes against concepts of efficient coding, which suggest that the system should dedicate more neural resources to stimuli that occur more often (25, 49, 60, 81, 82). However, these suggestions usually assume that the system has large capacity to process information and that there is no uncertainty in downstream circuits. This prediction also emerges in our task and model under the high resource-capacity assumption, but we argue that these efficient-coding assumptions hold only for cases in which organisms such as humans can process sensory information with high precision and have a perfect understanding of the task with no uncertainty about choice rules or related sources of noise in circuits dedicated to comparison, action, and learning. However, mice cannot follow human instructions and must learn task rules by trial and error, so it is reasonable to believe that behavioral variability in mice is largely impacted by downstream noise (46). Crucially, this counter-intuitive myopic behavior was evident in all the animals tested in this study, in line with the rational inattention framework. Although support for our theory from mice’s behavior is encouraging, we believe that these results might have important implications in human decision-making domains where the complexity and understanding of the task structure may be key to interpreting decision-making models. For instance, recent work has shown that a good deal of the variability in behavior of human participants in model-free and model-based learning processes might be rooted in task instructions (83). Thus, it is tempting to speculate that when task understanding is poor, agents may develop apparently deviant coding strategies that might be indeed efficient.

A key prediction of the rational inattention framework developed here is that information-processing strategies change endogenously as a function of stimulus-reward associations in a given environment, despite the fact that the physical environment is held constant across all contexts. Studies of efficient adaptation typically focus on scenarios in which he statistics of the physical environment change while keeping the reward contingencies constant. However, we argue that studying scenarios in which the statistics of the physical environment remain constant is also ethologically relevant, because physical environments are generally stable over long periods of time, whereas stimulus-reward associations may change more often. We found that mice’s characteristic levels of information-processing capacity in sensory and downstream circuits, enabled them to adaptively allocate their limited resources, which appeared to be related to improving learned associations that maximized utility. This led to myopic sensory encoding in various ways depending on the contextual reward-stimulus contingencies, as predicted by the theory.

Recent developments in the neurobiology of efficient information representation suggest that neural codes should not only be tuned by stimulus frequency statistics but also by the stimuli’s impact on downstream circuits and ultimately on behavior (29, 41, 42, 82, 84). In line with this intuition, there is evidence showing that early sensory systems represent not only information about physical sensory inputs but also non-sensory information depending on the requirements of a specific task and the behavioral relevance of the stimuli (55, 56, 85). Although we did not record the activity of sensory neurons, our work provides a formal justification for these intriguing observations, where information-processing resources endogenously adapt and reflect the organism’s needs. Given that noisy communication channels such as the brain always lose information during transmission, we argue that it is more efficient for the brain to adapt to the utility-maximizing rules of a particular environment at the earliest stages of sensory processing, an intuition that appears to be supported by recent imaging studies (56, 85).

In this work, we studied the possibility that a RL algorithm supported by operations that appear neurologically plausible could provide insights into how information-processing resources could be adaptively and optimally allocated through trial-by-trial experience. We leveraged recent discoveries suggesting that the brain can represent and update information about rewards in a distributional manner (63) and that functional remapping of task-relevant early sensory areas can be achieved with top-down feedback from decision-relevant circuits that encode prediction error signals and task rules (85). Interestingly, recent work revealed that dopamine neuron ensembles generate activity patterns that signal sensory-specific prediction errors (86), thus providing further support for our algorithmic architecture. Finally, adjustment of sensory gain is achieved via divisive normalization (87), an operation that has been related to the implementation of optimal attentional reallocation (67). We found that an algorithmic architecture of this kind allocates sensory resources in a context-dependent manner, similar to the patterns predicted by our rational inattention theory. Although in this model sensory resources are allocated dynamically and thus endogenously through trial-to-trial experience, in our current RL specification the parameters determining the learning rates are not endogenously estimated, but fitted to the data. A similar problem emerges with the original specification of distributional reinforcement learning (63): the agent needs to find a probability distribution consistent with the set of optimal updating operations, which requires a precise coordination in every time step of all neurons involved in RL computations. Although recent computational formulations provide hints how this problem could be tackled (88–90), it remains unclear how to connect these distributional RL computations to a biologically plausible algorithm that applies to arbitrary stimulus-reward association contexts such that reward expectation is maximized. This topic deserves attention and should be studied further.

Our computational model allowed us to investigate the role of adaptive regulation of the arousal system on sensory and nonsensory downstream imprecision. This approach revealed that nonspecific increase in behavioral sensitivity is associated with high levels of tonic arousal, but not directly related to sensory sensitivity. These results indicate that arousal systems may balance the costs associated with larger learning updates, which lead to high tonic arousal states and lower precision in downstream circuits while preserving sensory fidelity. This mechanism may have the benefit of leaving intact learning updates in sensory systems as a function of recently experienced sensory stimuli. Together, These results provide further support for the mechanisms of arousal-mediated adaptive gain control theory (91), indicating that some aspects of inattentive behavior might be indeed rational because the brain must operate under limited capacity constraints.

Additionally, we found that arousal systems carry reward expectancy information about specific visual cues that span the whole sensory space. An important implication of such neurocomputational signatures could be that agents track the utility distribution and volatility of the environment so that they can more accurately modulate prediction error signals. Interestingly, Recent work has shown that dopaminergic responses are amplified not only by the rarity of a reward received in a trial but also by its size (80). Thus, bottom-up noradrenergic reward expectation signals might be combined with top-down dopaminergic learning signals to efficiently guide reallocation of attentional resources in sensory systems. Thus, our computational modelling approach and behavioral paradigm implemented in rodents open the way to detailed investigations of the mechanisms underlying tight and essential interactions between arousal and dopaminergic systems.

Rodents have become an important model system in the study of decision behavior (44, 92–95). Here, we show that these organisms might be key to gaining a deeper understanding of the neurobiology underlying decision-making theories, which currently have a large impact in other disciplines such as medicine and economics. For instance, processes derived from rational inattention theories appear to be essential in guiding policy-making in both microeconomics and macroeconomic settings (96). In other fields, recent investigations have developed theories that attempt to provide normative accounts of complex neuropsychiatric conditions such as autism and schizophrenia, which have been characterized by deficits in performing optimal inference (97). However, these theories generally ignore the normative foundation that organisms must optimize behavioral processes in light of biological restrictions on information processing (98). Thus, the growing battery of molecular and imaging tools that is becoming available for use in rodents will enable a deeper understanding of the neurobiological underpinnings of limited cognition and apparently irrational decision behavior (45–47, 99). Thus, the corroboration of our theory using mice as a model organism opens the door to new directions that might be instrumental in the refinement and translation of these theories to applied settings in medicine, economics, and related social sciences.

## Methods

### Animal subjects

All animal experiments were performed in accordance with the Animal Welfare Ordinance (TSchV 455.1) of the Swiss Federal Food Safety and Veterinary Office, and approved by the Zurich Cantonal Veterinary Office. Subjects were adult (at least 8 weeks old) C57BL/6 mice (6 males, 1 female). Mice were kept on a reversed 12h/12h light-dark cycle, and all experiments were performed during the dark phase. For implantation of head plates, mice were anesthetized with isoflurane and placed into a stereotaxic frame (Kopf instruments). A custom printed headplate (Protolabs Inc) was fitted to ensure that the head was held at a natural angle in the behavioral apparatus. Before the experiments, the mice were allowed to recover from the surgery for at least 10 days.

### Behavioral setup and tasks

The specifics of the behavioural setup, as well as most of the parts used to construct it, are very similar to those developed in previous work (48), and widely adopted by the International Brain Laboratory (100).

The headfixed mice were trained to look at two gratings presented on the screen and pick the one that is more vertically oriented by turning the wheel with their fore-paws, which was coupled to the position of the two stimuli (Figure 2B).

Following recovery from surgery, the animals were put on a food restriction schedule (not dropping below 85% of initial weight). At the end of each training session the animals were weighted and additional amount of food was provided, dependent on the percentage of initial weight. For three days the animals were habituated to being headfixed and handled by the experimenter, with milkshake being delivered in random intervals during increasing periods of head-fixation. Then, mice were trained on the initial task, in which only a single stimulus appeared on either left or right side of the screen, which they had to move to the center of the screen (response location), in order to obtain a milkshake reward. After they mastered the simple slection task, the second grating was introduced (more horizontal/distractor), initially at low contrast, but progressively increasing contrast across sessions as the animal learned. After the second grating was completely introduced (contrast = 1) and session performance was above 60%, we started varying the contrasts of the two gratings independently across 3 levels (0.3, 0.6 and 1). Importantly, during the entire training procedure, we kept the orientation stimuli *θ* were drawn from the prior distribution *π*(*θ*) (see main text and Figure 1A). Once the animals reached good performance on the task (>60% across difficulties), we introduced different mappings of reward to orientation (Ω). In each of these mappings the reward size obtained for correctly picking the more vertical orientation was dependent on that orientation. We then ran each animal in each of the reward environments for at least 20 sessions. Here we studied adaptation to particular environment Ω after 7 sessions of experience. Considering longer adaptation periods does not affect the main conclusions of our work.

To keep the volumes of milkshake constant the opening times of click valves (NResearch 161K011) were calibrated, before each day of training, to deliver a constant volume of milkshake. This was achieved by opening the valves repeatedly for different durations and subsequently fitting a curve, describing the relation between opening time and ml of delivered milkshake. Milshake amounts in μL were mapped to the correct-side angle, dependent on the reward mapping environment Ω. In the 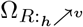 environment, the reward mapping was a linear increase from horizontal (0 deg, 1 μL) to vertical (90 deg, 8 μL). The reward mapping for the 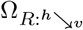 environments was exactly the opposite, with horizontal grating yielding 8 μL and vertical 1 μL. In the Ω_*R*:*h*⇄*υ*_ environment all angles yielded 5 μL. The illustration of the reward mappings for each environment Ω is shown in Figure 1E, and the exact amounts are shown in Supplementary Table 1.

The stimuli were Gabor patches each spanning 15 visual degrees and with the phase of the grating randomised between trials. Decision was considered made when one of the gratings was brought to the middle of the screen. Correct choice yielded a drop of strawberry milkshake, and incorrect led to white noise being played for 0.5s and a 2s timeout before next trial (in addition to the ITI of 1.5 – 2.5 sec). To counteract bias, where animal might be content with just turning the wheel one way and getting 50% of rewards, incorrect trials were repeated (correct side was kept the same, but orientations resampled), but data from these trials were not used in the final analysis. To check that this did not influence the results, two animals were successfully trained also without repeating incorrect trials. For all experimental conditions, the orientations of the two gratings were picked from a manually set prior distribution of orientations (Figure 1A), with the difference between the two ranging from 20 to 90 degrees, in increments of 10. In a small percentage of trials, the two orientations were the same, in which case a random side choice delivered reward. For half a second after stimulus presentation the stimulus position was not coupled to wheel rotation, but the animal was not punished for turning it at this time (Figure 1C). Signal tone (12kHz), played for 0.2s cued the end of wheel-uncoupled phase and the animal could make the response.

### Pupil size analysis

In order to track the pupil and licking, we filmed the animal from the side, illuminated by infrared light, using the FLIR Blackfly^®^ camera. Pupil edges and licking responses were extracted using DeepLabCut (101) and eye diameter was calculated from the obtained videos. Trials were synced to video using a small white square in the corner of the screen that appeared at stimulus onset. The area of the pupil was calculated from the points tracked by DeepLabCut, and the trace for each daily session was lowpass filtered with a cutoff frequency of 20Hz to remove noise. Pupil size was then z-scored within session. Pupil size was binned into 0.2 second bins, with the binning window moving by 0.1 second, further smoothing the data. All multiple linear regressions studied here were ran on binned time-points with pupil size as dependent variable. Factors in the regression included: absolute angle difference, contrast sum, reaction time, correct side orientation, correctness on a trial, correctness on the previous trial and binned licking. Furthermore, we split the data into correct and incorrect trials, and ran the same regression, but excluding the trial correctness parameter. We also measured phasic pupil responses by subtracting the baseline 0.5 seconds before stimulus onset from each trial pupil size. To investigate the arousal responses based on reward expectancy, we ran a bin-time regression to predict the phasic pupil response, same as for the non-baseline corrected data.

### Descriptive behavioral models

Here we studied three descriptive models, each with different levels of complexity, which allowed us to evaluate basic behavioral signatures in all three different reward mapping conditions. This family of models do not explicitly use information about the statistics of the environment and condition-dependent behavioral goals.

On each trial, mice face two alternatives *l* and *r* with corresponding orientations *θ_l_* and *θ_r_*, which are shown on the left and right side of the screen, respectively (Figure 1B). The goal of the mice in all three stimulus-reward environments Ω is to select the alternative that is closer to 90°. In the description of all models (unless otherwise specified), we assume that the orientations are mapped to an abstract “verticality” space from 0° to 90° (e.g., *θ* = 170° is mapped to *θ* = 10°). In descriptive model 1 (DM1), the probability of choosing alternative *l* in trial *n* is given by

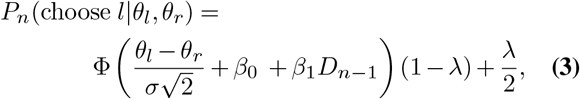

where Φ() is the normal cumulative density function, *σ* represents the degree of sensory noise in the representation of *θ* (which in DM1 model is fixed for all orientations), *β*_0_ captures potential side biases, *β*_1_ captures biases caused by the decision *D* in the last trial (*n* − 1), and *λ* captures potential lapse rates (in order to simplify notation, in the remainder of the model descriptions we will drop the trial indexes *n*).

In descriptive model 2 (DM2), we studied the possibility that different levels of contrast are associated to different levels of sensory noise. Therefore, DM2 is given by

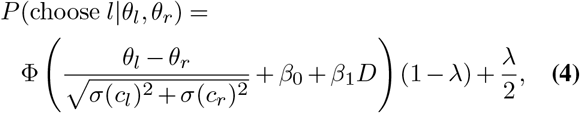

where in this case the noise level of each alternative is a function of contrast *c*. In this model we fit three different noise levels for the different levels of contrast used in this study.

Finally, in descriptive model 3 (DM3), we investigated the possibility of “risk seeking/aversion” depending on the level of sensory uncertainty of each alternative induced by the different levels of contrast.

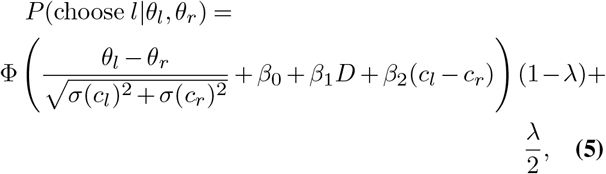

where in this case *β*_2_ determines the strength of “risk seeking/aversion” (with negative values corresponding to apparent “risk aversion”).

### Rational inattention model

We studied the possibility that mice behave according to the rules of optimal statistical inference in estimating the orientation of each choice alternative. Crucially, here we take into consideration the fact that organisms have limited capacity to process information, and therefore, neural systems must allocate these limited resources according to each reward-mapping context in order to maximize the amount of reward consumed over the course of many trials and days.

We assumed that for a given input *θ*_0_ the mouse makes an internal measurement *m*(*θ*_0_) which is corrupted by sensory noise. Crucially, the level of sensory noise will depend on how many resources the system dedicates to the measured orientation (described in detail below). Each time orientation *θ*_0_ is presented to the observer, it results in different measurements *m_i_* described with a conditional probability function *p*(*m_i_* | *θ*). On each trial *i*, the observer computes a posterior distribution *p*(*θ* | *m_i_*) by combining the physical environmental prior distribution *π*(*θ*) with the likelihood of the measurement *p*(*m_i_* | *θ*). Then the mouse applies a decoding rule in order to obtain a posterior estimate 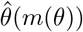. Here we assume that mice compute the expected value of the posterior distribution: *E*[*p*(*θ* | *m*)]. We assume that on each trial, the mouse independently estimates 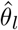 and 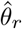 for the input orientations *θ_l_* and *θ_r_*, respectively.

The measurement *m* follows a von Mises distribution

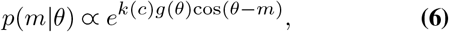

where the precision of the measurement is determined by two multiplicative factors: (1) *k*(*c*) which is function of the contrast level for a given orientation in a particular trial (see Eq. 7 below), and (2) *g*(*θ*), a limited-resource gain function. Here *g* acts as a gain control mechanism that regulates how many resources the system allocates to a particular segment of the orientation space.

Activity of visual neurons as a function of contrast has been well described by the following function

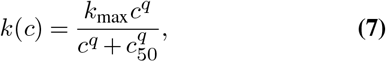

where *k*_max_ is the maximum activity of the neurons, and *q* and *c*_50_ specify the slope and semi-saturation point of the contrast response function (102). Given that the variability of cortical neurons approximates a Poisson distribution, one can assume that the relative variability in the measurement *σ*(*c*) decreases in inverse proportion to the square-root of the cortical activity. Because the precision in the von Mises distribution can be defined as *k* = 1/*σ*^2^, neural precision as a function of contrast can be described by equation 7.

If we constrain *g*(*θ*) > 0 for all *θ* and to be a normalized function (we formally explain how *g* is estimated below), and assume a cost function *K* that provides information about the metabolic resources employed to encode visual information, one can formulate an optimization problem in order to find the optimal allocation of resources in the orientation space for: (i) a given physical environmental prior *π*(*θ*), (ii) a given contrast response function *k*(*c*), (iii) reward outcomes associated to decision-outcome rules in a given context or environment Ω, and (iv) downstream noise *σ*_late_ that is not related to sensory encoding. Formally, the goal is to find a resource allocation *g*\*, and the maximum allowed activity *k*_max_ (with an underlying contrast response function, Eq. 7) such that

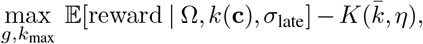

where **c** is the set of contrasts that the animal experiences. We model the metabolic cost *K* as a linear function of the average precision 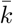 that the agent invests on solving the inference problem, thus 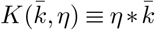, where *η* > 0 indicates how much the metabolic cost scales with average precision 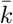 defined as

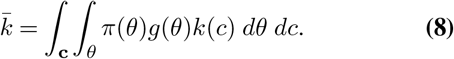

We could have also considered more complex non-linear relationships between metabolic costs and neural activity. However, the main conclusions of our work are not affected by the exact specification of the cost function, and it has been demonstrated that the linear relationship is a reasonable assumption (59). We also note that we have not included costs associated to downstream precision 1/*σ*_late_. However, if we assume that *σ*_late_ influences choices (and therefore reward expectation in 2), including such cost in *K* as an additive factor does affect the estimation of the optimal allocation of resources *g*.

Here we do not employ information transmission as optimization criteria as classically assumed in the rational inattention literature, which is defined as follows:

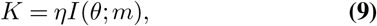

where *I*(*θ*; *m*) denotes the mutual information, which measures the expected reduction in uncertainty after observing a signal *m*

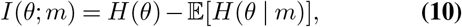

where *H*(*θ*) denotes the entropy of the prior distribution *π*(*θ*) and the second term measures the expected reduction in uncertainty after observing the signal. While this cost function has several benefits (such as mathematical tractability for relatively simple problems), it also has been pointed out that specifying attention costs to be linear in entropy reduction might be problematic in particular when applied in problems of sensory perception (43). In any event, for the case of sensory perception, and in our experimental paradigm, we argue that it is more appropriate to assign costs to the amount of energy invested in generating neural activity as defined above. Nevertheless, we show that the two costs functions generate similar qualitative predictions in the resource allocation function *g* (Figure S3), and also study the relationship between expected precision and expected uncertainty reduction (Figure S4).

The next step is to specify the optimization problem. For instance, in the “constant reward for correct decisions” condition (i.e., environment Ω_*R*:*h*⇄*υ*_), the agent aims at minimizing the expected probability of errors plus its associated metabolic costs

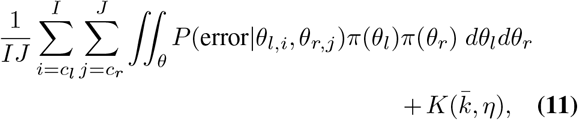

where indexes *i* and *j* reflect the the different levels of contrast (three in our experiments) applied to the left and right orientation inputs, respectively. Here, we still need to define the probability of making an erroneous response *P* (error). In our observer model, a source of decision errors is caused by variability in the estimates 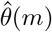 due to measurement errors *m*, thus resulting in a resulting in a conditional probability of estimated orientation given the true stimulus orientation 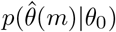 (in order to simplify notation, we leave occasionally the dependence on *m*). Here, we assume that the distribution of estimators 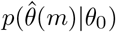 is Gaussian and therefore it will be convenient to compute its expected value 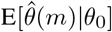 and variance 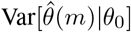, which are given by

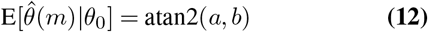

and

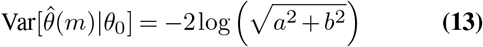

respectively, with

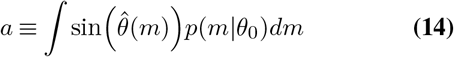

and

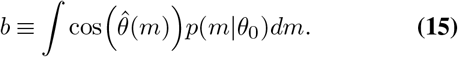

As in the descriptive models, here we assume that the orientation estimations are mapped to a “verticality” space, and therefore the probability that the mouse selects *θ_l_* is given by

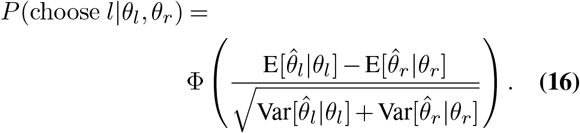

In addition to sensory precision in the coding of orientation *k*(*c*)*g*(*θ*) (see Eq. 1), we also account for late noise in the decision stage (that is, post-decoding noise), which may capture any unspecific forms of downstream noise occurring during the response process that are unrelated to the estimation of orientation per se. We assume this late noise to be unbiased and Gaussian distributed 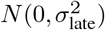; therefore, it can be easily added to our model as follows

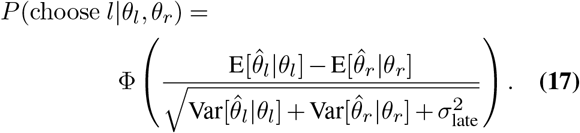

Hence, the probability of an erroneous decision *P*(error) in Eq. 11 can be defined as follows:

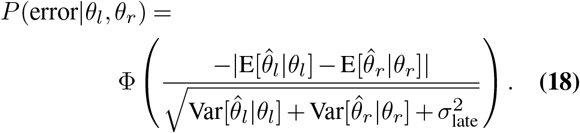

With these definitions, we can formulate the optimization problem for the remaining reward mapping conditions. For the environment Ω in which the amount of received reward *R* for correct responses is linearly mapped to the degree of “verticality” where horizontal (*h*) orientations are mapped to the smallest reward and vertical (*υ*) orientations are mapped to the highest reward (environment denoted 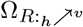), and receive no reward for incorrect responses, the goal is to find the allocation of resources that minimizes the following expression

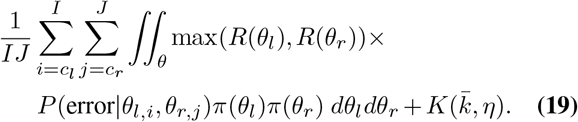

For the environment Ω in which the amount of received reward *R* for correct responses is linearly mapped to the degree of “verticality” where horizontal (*h*) orientations are mapped to the highest reward and vertical (*υ*) orientations are mapped to the smallest reward (environment denoted 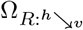), and receive no reward for incorrect responses, the goal is to find the resource the resource allocation function *g*(*θ*) that minimizes the following expression

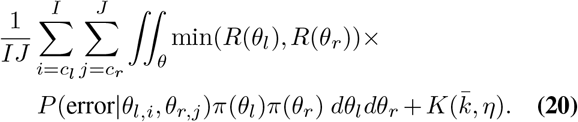

Recall that in all cases the decision rule is to choose the orientation that is more vertical. Therefore, in the 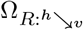 environment we are actually asking the mice to choose the orientation that is mapped to the smallest reward, however, in this environment choosing the smallest orientation on any given trial is actually the decision that guarantees reward receipt.

The next step is to describe a methodology that allowed us to find potential solutions in our resource allocation problem. Our approach is to estimate the minimum achievable reward loss (plus costs) in a given environment Ω, by finding the minimum achievable value over a flexible parametric family of possible functions *g*(*θ*), with the following properties

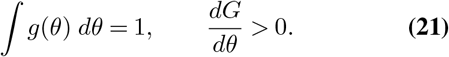

Here we assume a finite-order polynomial function *G*(*θ*) consistent with the property that *g*(*θ*) = *G*(*θ*)′. This requires *G*(0) = 0 and *G*(1) = 1, and therefore *G*(*θ*) can be written in the form:

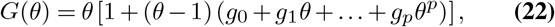

with the properties described in Eq. 21, where **g**_*p*_ = {*g*_0_, … , *g_p_*} is a set of parameters over which we optimize. Note that for a large enough value of *p*, any smooth function can be well approximated by a member of this family. Also note that for the case *g* = 0, … , *g_p_* = 0, the resource allocation problem is given by a rule that assigns equal ammount of resources to all orientations *θ*.

In this study we use a parameter vector **g**_*p*_ of order *p* = 2 (we found that using higher order did not significantly improved the optimal solutions). Given the symmetry of the prior distribution relative to *θ* = *π*, we estimate an initial 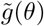 for the space *θ* ∈ [0, *π*] and then defined 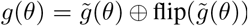, where the operator ⨁ denotes concatenation and the operator flip() denotes vector reversal.

### Rational inattention model predictions

We used the empirical distribution and input stimuli as well as the distribution of rewards (i.e., drops of milkshake) used in our experiments in order to derive predictions of the resource allocation function *g*(*θ*), the maximum activity allowed activity *k*_max_, that minimized reward loss (plus its associated sensory precision cost *K*). Given that we were interested in studying how resource allocation is potentially influenced by different levels of down-stream noise *σ*_late_, we found the optimal parameter vector **g**_*p*_ and *k*_max_ for different combinations of *η* and *σ*_late_ (Figure 2 and Figures S2,3).

### Applying the rational inattention model to empirical data

In order to fit the inference model to the empirical data in a way that was comparable to the full descriptive model (DM3, see above) we used

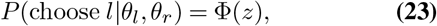

with

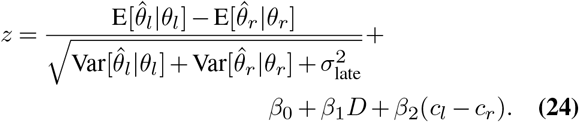

For all models, we are interested in finding the set of parameters ***β***, *c^q^*, 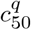 and *σ*_late_. For the endogenous model, we were in addition interested in finding *η*. For the exogenous model, we also found **g** and *k*_max_. Notably, the endogenous model has the same number of parameters as the most complex descriptive model (DM3). Additionally, note that we dropped the lapse rate parameter *λ* from the rational inattention model. We reasoned that the rational inattention model rationalizes the apparent lapse rates that emerge in the descriptive models. In fact, including *λ* as a parameter in our model led to an estimated valued *λ* ≈ 0 for all mice, thus confirming our intuition.

In order study whether the reallocation of limited resources is necessary to explain the data, we also considered a model where **g**= {0, 0, 0}, which corresponds to a model where mice allocate their resources equally across the whole sensory space.

### Learning model

We investigated whether a reinforcement learning (RL) model that incorporates the same coding constraints of the ideal observer model would generate similar performance. That is, in the learning model we do not directly find the parameters **g** of the gain coding function *g*(*θ*), but this function is continuously updated based on trial-to-trial experience via RL (Figure 3A).

Assume that in any given trial at time *t* a mouse chooses alternative 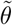. After this decision, the mouse receives reward *R_t_* that depends on the chosen option 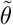 according to decision-outcome rules of environment Ω (Figure 1C). In addition, here we assume that mice evaluate their degree of confidence *C_t_* based on the decision in trial *t*. Here, we define confidence based on the statistical definition of confidence (103), that is, the probability that the chosen option is correct given the chosen option: 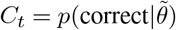. Information about *R_t_* and *C_t_* is then used to update a reward distribution vector 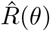 (mapping orientation stimuli to reward) using the reinforcement learning rule

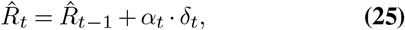

where, *α* is defined as the learning rate, which can take one of two values that depend on positive and negative prediction errors

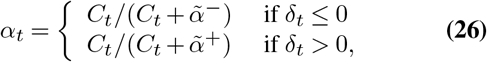

with 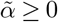. This implies that if 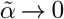, then the learning rate *α* → 1 and confidence *C_t_* have little influence. On the other hand if 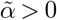 then confidence starts to have more influence in the learning rate, with higher values of confidence having a stronger influence in the update rule. We define the prediction error vector

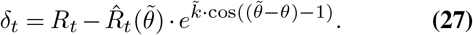

This definition implies that the reward distribution vector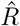 is updated according to the location of the chosen stimulus 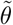 smoothed by parameter 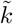, and modulated by learning rate *α_t_*. Finally, the reward distribution 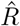 is transformed to the gain function *g* via a normalization operation

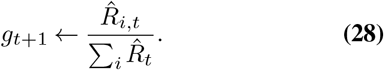

Hence, this RL model has only three free parameters that are fitted to the observed data: 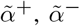, and 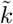. Vector 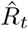 was discretized in steps of 2° from 0° to 178° in the orientation space.

### Model fits

For a given model *M*, we denote its set of parameters by a vector **Ψ**. The goals is to find the combination **Ψ** that maximized the probability of all of a single subject’s responses given the presented stimuli and the parameters. In this case, the log of the parameter likelihood function is

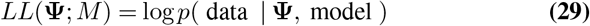

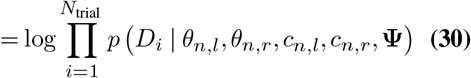

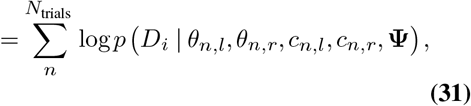

where *θ* and *c* are the orientation and contrast inputs for the left (*l*) and right (*r*) alternative, respectively, and *D_i_* is the mouse’s response on trial *i*. The implementation of the likelihood function was implemented using the mle2 function in the bbmle library (104) implemented in the software package R (105). We typically performed an initial stage with 2,000 randomly chosen initial parameter combinations. For each model *M*, we repeated this procedure three times, leading to log likelihood values that were typically within one point. We are therefore reasonably confident that we found the global maxima for each model. As model comparison method, we use the Akaike information criterion (AIC) (106).

### Poisson neural model

We investigated if a more biologically relevant model of V1 function would also be able to capture characteristics of the rational inattention codes observed in the algorithmic model. An advantage of this implementational approach is that we can relate the metabolic cost directly to the expected number of spikes that the system generates to solve the decision problem.

The neural population consisted of Poisson neurons, each generating spikes *r* independently for a given input stimulus *θ* with probability

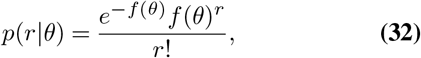

The tuning functions *f* of the neurons follow a bell-shaped activation pattern of the form

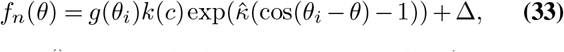

where *g*() is a multiplicative gain partially determining the distribution of coding resources through the orientation space, *k*(*c*) is the multiplicative gain that the determines firing rate as a function of contrast, *θ_i_* is the preferred orientation of neuron i, and Δ the base firing rate of a neuron at rest. 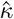 controls the width of the tuning curve. The tuning curve width 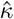 was estimated from the same study that we used to derive the prior distribution *π*(*θ*) (47) (see Results section in main text and Figure S1).

Based on this specification, the log-likelihood function of population vector activity 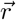 can written as

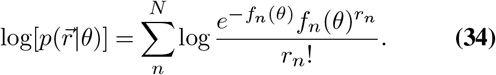

To derive a posterior estimate 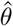, we assumed the Bayesian least squares estimate (BLS). The BLS estimator can be approximated with discrete sums,

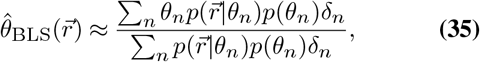

in which *θ_n_* is a discrete set of stimulus values and *δ_n_* is the spacing between adjacent values. When *θ_n_* (the preferred stimuli of the neurons), are distributed in the stimulus space proportional to the prior distribution *θ_i_*, then 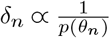 (60, 71), thus, the BLS estimator can be written as

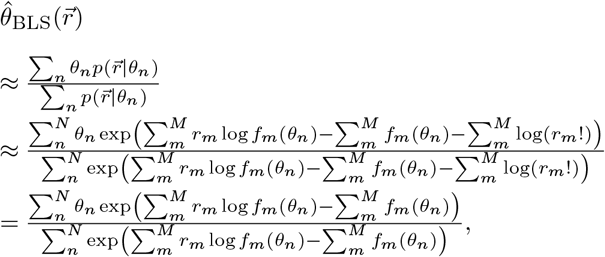

where in the last step, the last sum inside the exponential does not depend on the stimulus and can be dropped from the numerator and denominator.

Applying this model to our decision task, we assume that for two inputs *θ_l_* and *θ_r_* in any given trial, the model generates population activity 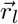 and 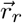 independently for each input, which leads to the computation of estimates 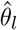 and 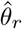. Then a choice is made based on the decision rule of our task.

The goal of the neural network to find a balance between model performance (i.e., reward intake) and the metabolic costs associated to neural activity in in the network. More formally we need to find the optimal resource allocation *g* and a maximum activity *r*_max_ such that

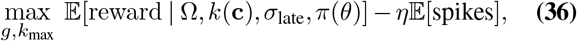

where in this case *η* corresponds to unit cost per spike generated by the network.

## ACKNOWLEDGEMENTS

We thank Anne Urai for valuable discussions during the setup of the mice experiments. We would also like to thank Peter D’Almeida and Tom Christen for assistance with animal training. This work was supported by a European Research Council (ERC) starting grant (ENTRAINER) to R.P. This project has received funding from the European Research Council (ERC) under the European Union’s Horizon 2020 research and innovation programme (grant agreement No. 758604), and ETH Zurich (D.B.).

## AUTHOR CONTRIBUTIONS

N.G. built the experimental setups, piloted the development and coding of the experimental paradigm, performed all surgeries, trained the animals, and carried out data analyses. J.B. contributed to the experimental code, trained the animals, and implemented computational models. R.P. conceived the original idea of the study, and contributed to data analyses. All authors contributed to conceptual development of the experimental paradigm, discussed the results and wrote the paper. D.B. and R.P. raised funding and jointly supervised the study.

---

**Supplementary Figure 1.**
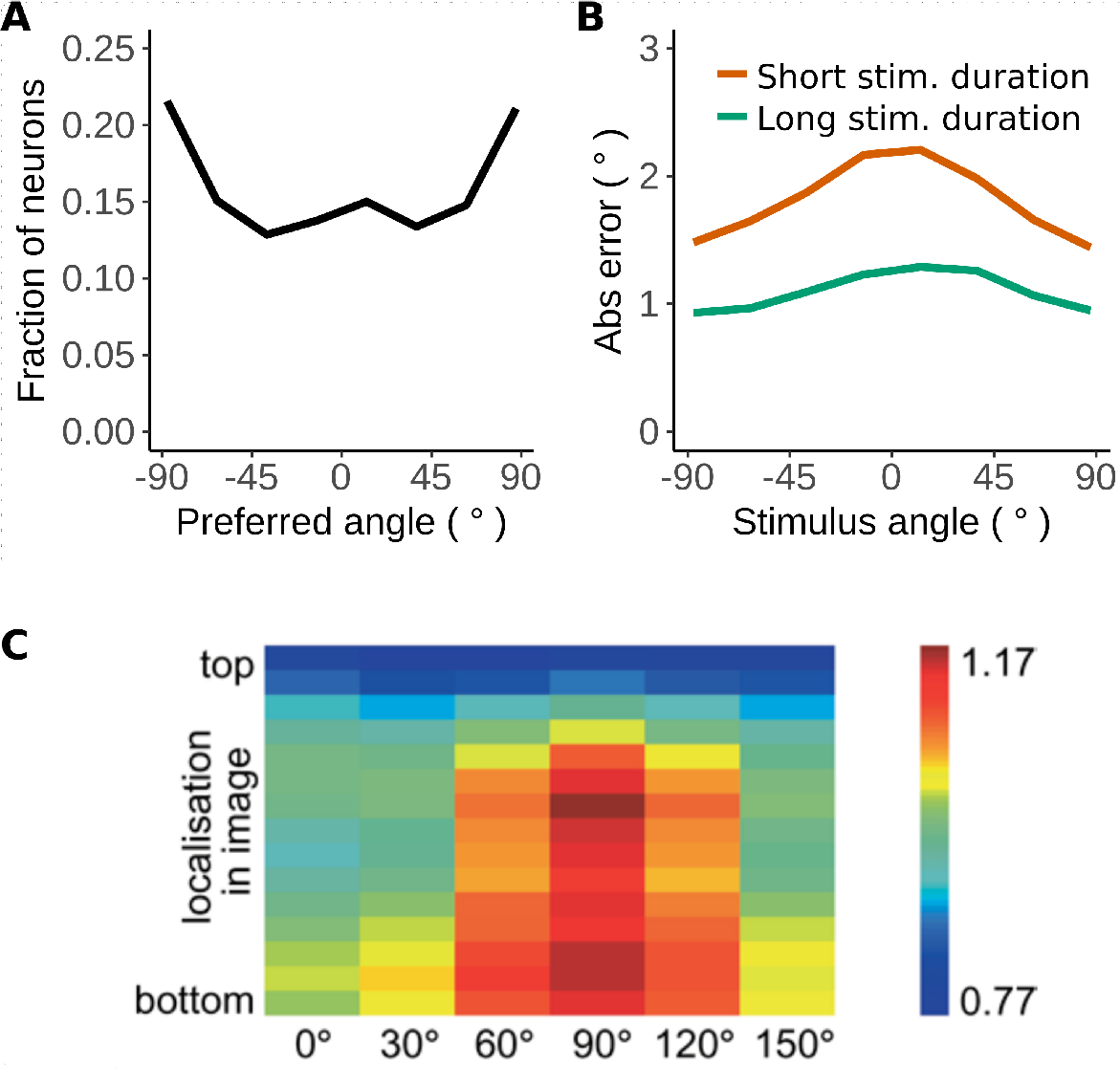
Mouse V1 has more coding resources at the horizontal angles. Carsen et al. recorded multi-plane two-photon calcium images from the primary visual cortex of awake mice (47). Stimuli were static gratings rotated at random orientations. **A)** Each neuron has a tuning curve with a strongest response to a preferred orientation. The distribution of preferred orientations across cells shows more neurons prefer horizontal angles. **B)** Carsen et al. used a linear decoder to estimate the stimulus orientation from neuronal activity. They found that coding error is higher for the vertical angles. When the stimulus duration was shortened (100 ms (red) vs. 750 ms (green)) the decoding error of the linear decoder goes up. The results presented in A) and B) are indicative that mice are more exposed to horizontal than to vertical orientations in the natural world. To simulate the natural exposure over orientations the prior distribution *π*(*θ*) used in our behavioral task is also higher for horizontal than vertical orientations. This allows us to study the effects of changing reward-stimulus contingencies on the coding strategies, in an environment that keeps the natural prior neural codes untouched. **C)** Betsch et al. mounted a camera on top of the head of 4 cats to record natural stimulus videos of freely behaving cats (107). The mean wavelet amplitude of the recorded videos is shown for 6 orientations on the x-axis. The y-axis represents the height in the recorded image. Most of the visual field is dominated by horizontal orientations. Although this study was performed using cats instead of mice, it is interesting to observe that horizontal orientations dominate the natural environment of this species.

**Supplementary Figure 2.**
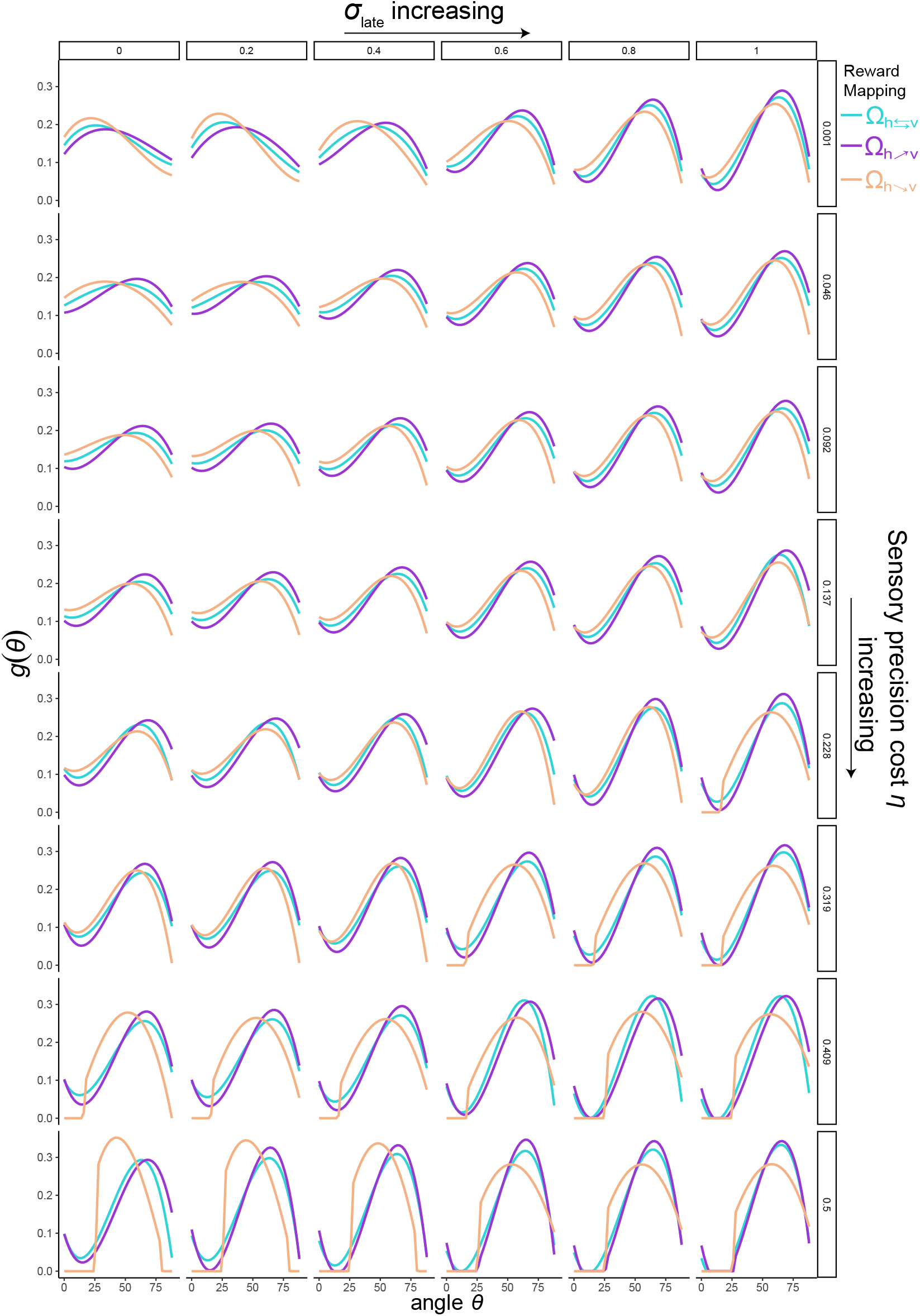
Optimal *g*(*θ*) depends on precision costs and noise. The way in which the system should invest in sensory encoding for a particular portion of the stimulus space depends on both precision costs *η* (increasing from top to bottom) and late noise *σ*_late_ (increasing from left to right). An interesting property of the rational inattention model is that as both *η* and *σ*_late_ increase, the less resources the system invests on regions of the stimulus space with higher density. This demonstrates important differences between solutions of efficient coding that that assume low levels of noise in the encoder (e.g., top left panel) and solutions where the low-noise regime is dropped.

**Supplementary Figure 3.**
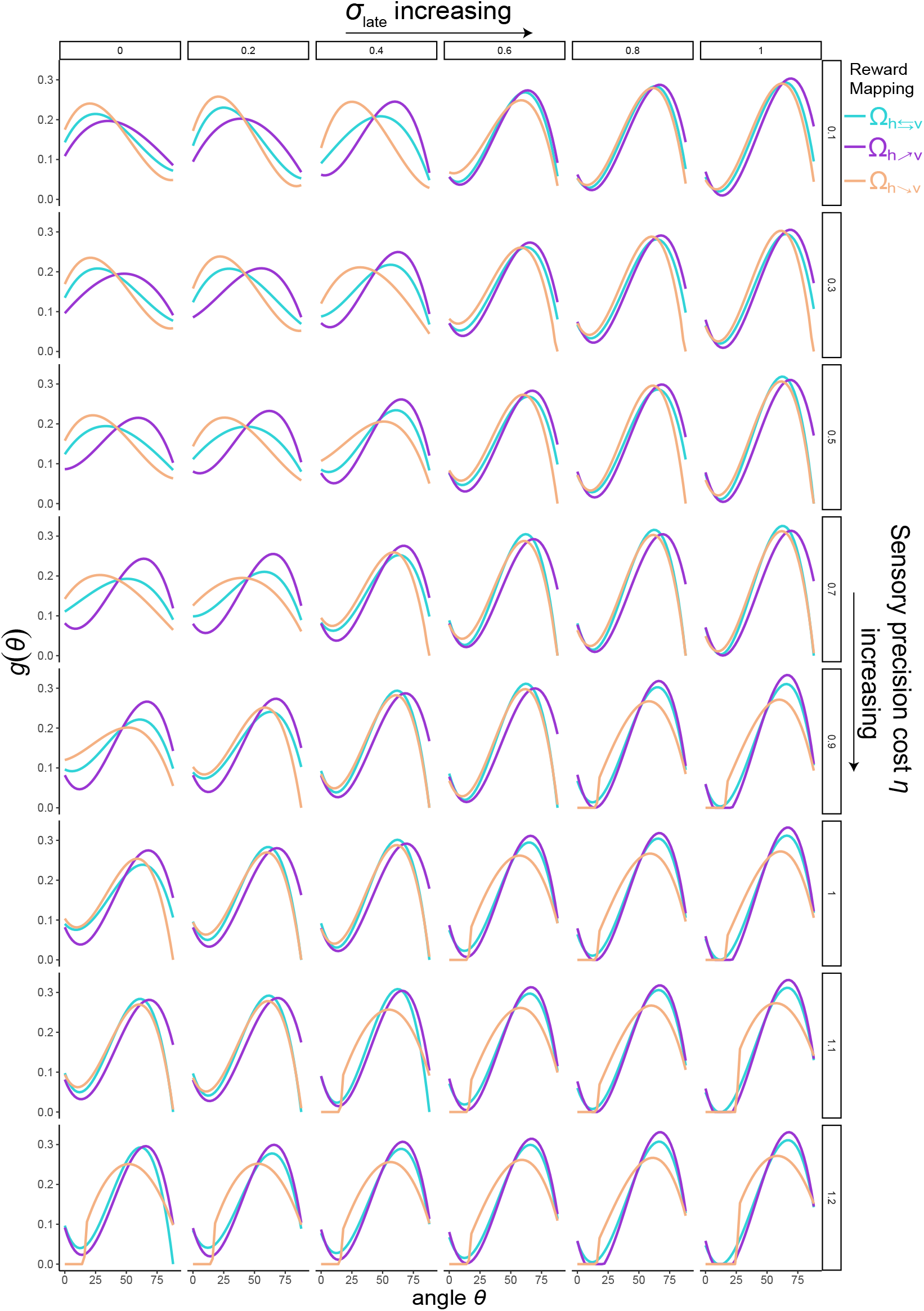
Optimal *g*(*θ*) using entropy as cost function. This figure shows similar information to Supplementary Figure 2, with the difference that now the cost function is the expected reduction in entropy 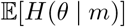. While there are slight differences in the solutions for *g*(*θ*), these are in general qualitatively similar to the solutions for using precision as cost function. The reason is that both cost functions increase with the use of more resources. However, the relationship between expected precision and entropy reduction is not linear (see Supplementary Figure 4).

**Supplementary Figure 4.**
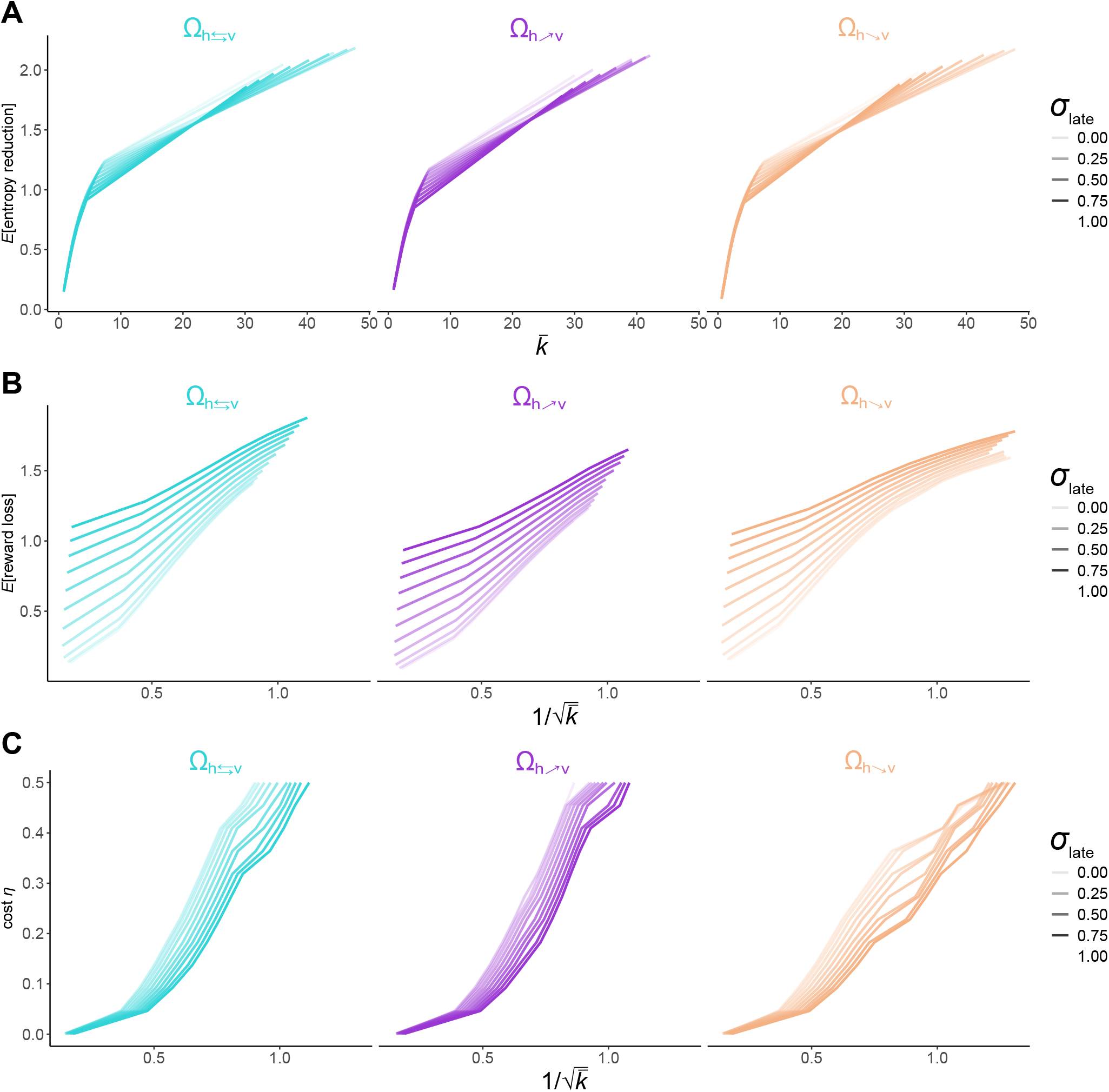
Relationship between average precision, entropy reduction, reward loss, and cost. **A)** Relationship between average precision 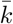 and entropy reduction 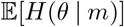 for the rational inattention model optimized for a linear precision cost for each environment Ω (left, middle, and right panels), and for different levels of late noise *σ*_late_ (different levels of line transparency). The results of this analysis show that the relationship between these two metrics is non-linear, showing signatures of concavity. This suggests that optimizing based information transmission investment is more liberal than optimizing based on precision investment. **B)** Relationship between the inverted squared root of average precision 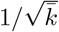 and expected reward loss for the rational inattention model optimized for linear precision cost for each environment Ω (left, middle, and right panels), and for different levels of late noise *σ*_late_ (different levels of line transparency). As expected, the higher 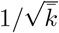 and *σ*_late_, the higher the expected reward loss. **C)** Relationship between 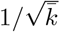 and linear precision cost for linear precision cost *η* for each environment Ω (left, middle, and right panels), and for different levels of late noise *σ*_late_ (different levels of line transparency). As expected, the higher 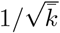 (i.e., smaller invested precision) the higher cost. Also note that for a fixed cost level, the smaller the level of late noise *σ*_late_, the lower the precision that the organism/system should invest.

**Supplementary Figure 5.**
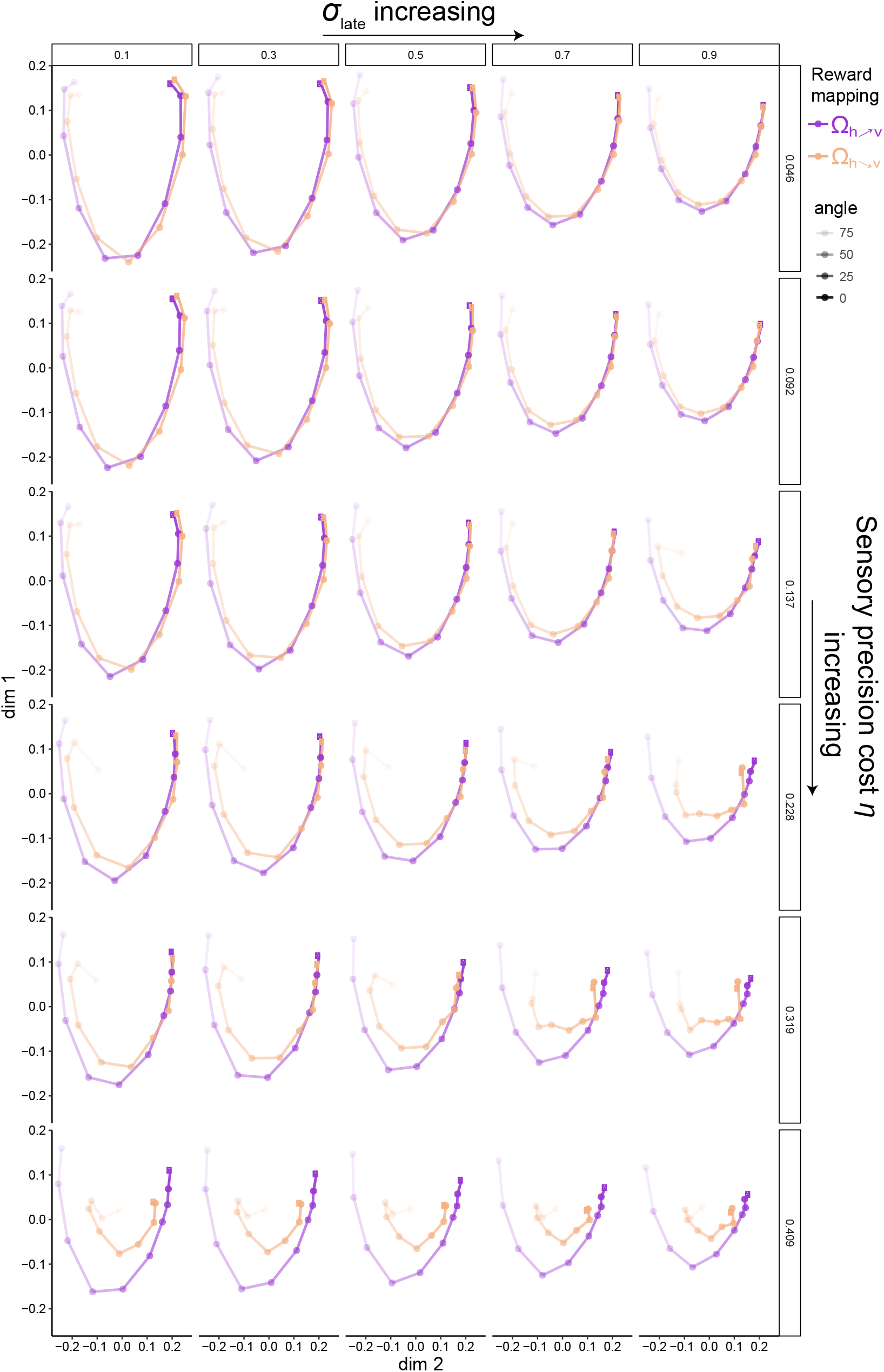
Geometric analyses of psychophysical performance of the rational inattention model allowing sensory resources *g* to be adaptive across the sensory space. In order to have a better overview of predicted discriminability performance across all pairwise combinations of input stimuli as a function of precision cost *η* and downstream noise *σ*_late_, we implemented a multidimensional scaling (MDS) analysis. For clarity in the presentation of these analyses, we considered only environments 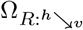 and 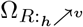, which are diametrically orthogonal in terms of stimulus-reward mappings. The introduction of MDS analyses is appealing because it provides an intuitive interpretation of behavioral performance. Here, each dot represents an input stimulus (where each angle in is represented with a different transparency level). The distance between a pair of nodes int the manifold then represents degree of discriminability between the two angles. In our task the MDS various important features of psychophysical performance: (i) The higher the sensory precision cost *η* and downstream noise *σ*_late_ the shorter the distance between dots, thus indicating lower levels of overall discriminability. (ii) In enviroment 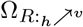 discriminability at horizontal angles is smaller relative to environment 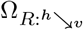, and vice-versa for vertical angles. (iii) For a given level of sensory precision cost *η* and downstream noise *σ*_late_ psychophysical performance is better in environment 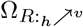 relative to 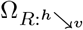. This prediction emerges in the rational inattention framework because of the resource constraints and inference process considered here. For a given level of sensory and decision precision, reward expectation in environment 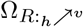 is higher. Thus our rational inattention model generates a set of testable predictions that can be verified with the empirical data.

**Supplementary Figure 6.**
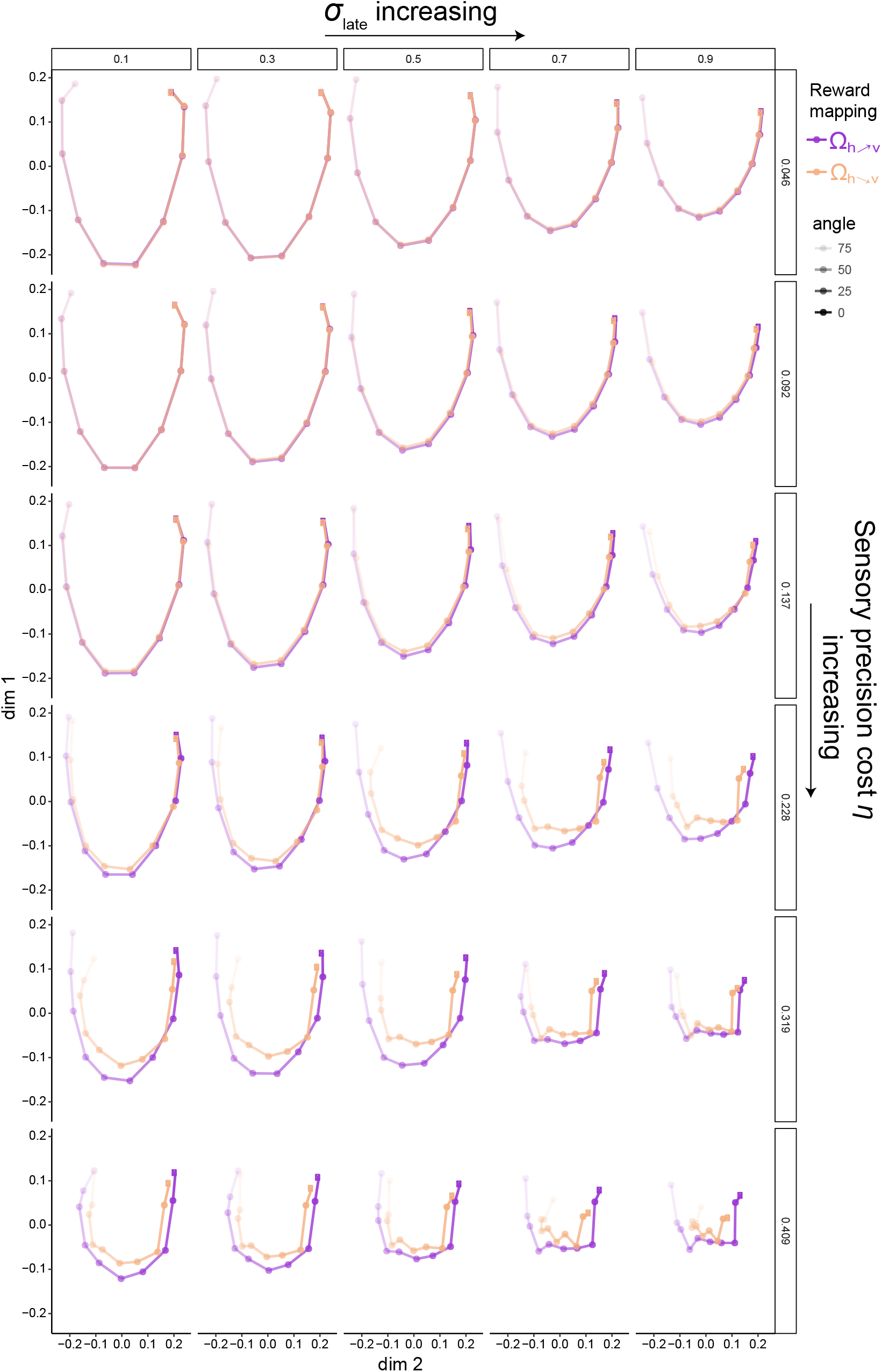
Geometric analyses of psychophysical performance of the rational inattention model assuming uniform resource allocation *g* across the sensory space. Here con conducted MDS analyses in the same way as conducted in Supplementary Figure 5, but this time assuming uniform resource allocation *g* across the sensory space. While for a given level of sensory precision cost *η* and downstream noise *σ*_late_ psychophysical performance is better in environment 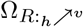 relative to 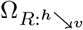, and thus similar to the prediction presented in Supplementary Figure 5 (a general prediction of the rational inattention framework), the degree discriminabilty is relatively constant across the sensory space. Additionally, for the same levels of sensory precision cost *η* and downstream noise *σ*_late_, the degree of discriminability is larger for the variable gain *g* rational inattention model (Supplementary Figure 5) relative to the uniform gain *g* model (this figure). This is qualitatively evident by comparing the overall distance between the nodes of the manifold across Figures S5 and S6. Thus the two types of rational inattention models provide distinct qualitative features that can be qualitatively compared against empirical data.

**Supplementary Figure 7.**
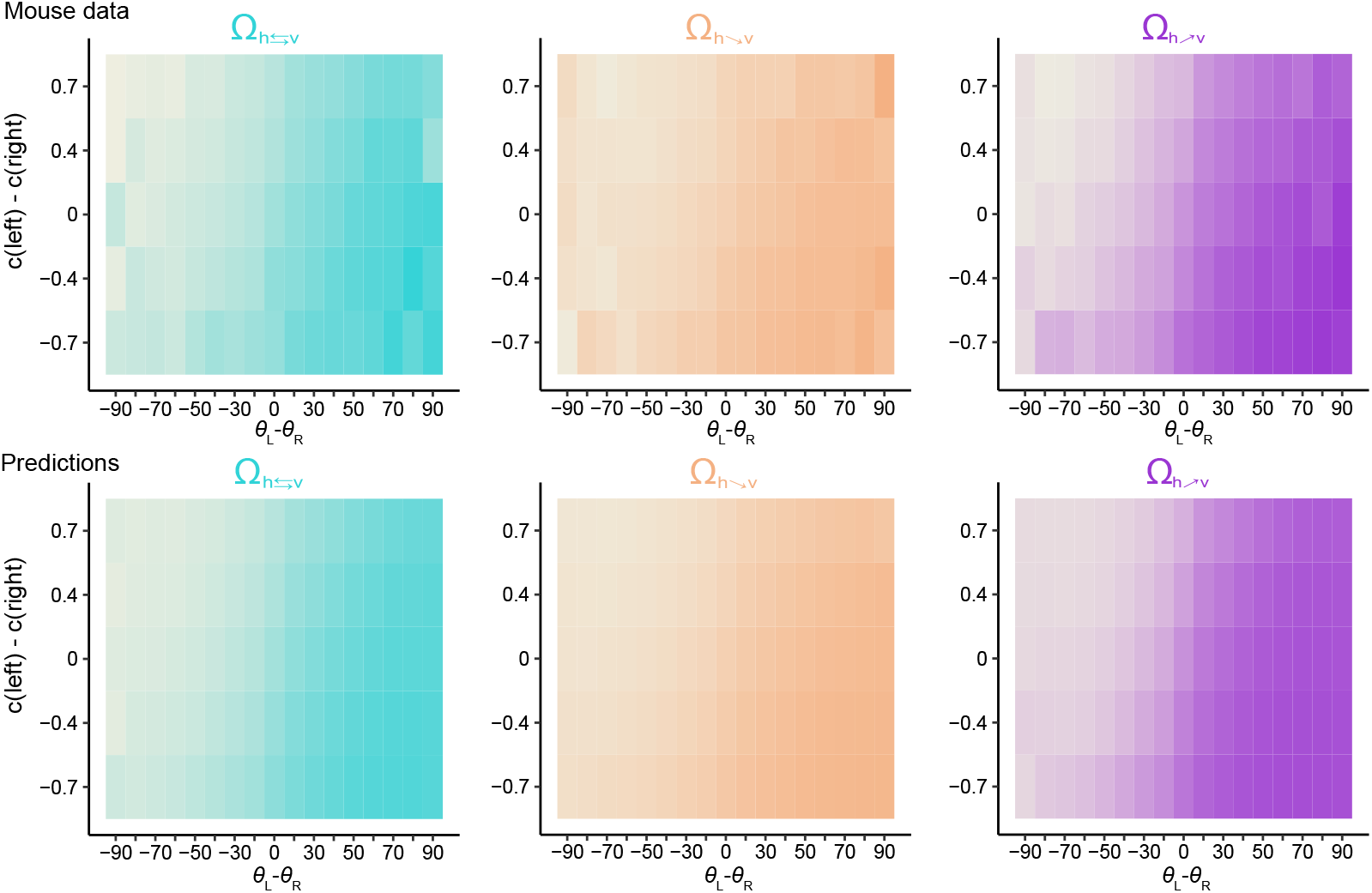
Mouse data and rational inattentive model predictions for difference in contrast levels. Heatplots for each reward environment (denoted on top of heatplot and color coded) show probability of choosing the left stimulus at each contrast difference in the study vs. the angle difference (trial difficulty). Top three plots show real mouse data and bottom three show predictions of the static model. The contrast differences between two stimuli that were 0.4 and 0.3 were grouped together for the purpose of this plot, and are marked 0.4 on the y axis. Preference for higher contrast side stimuli as a function of difficulty, which could be interpreted as risk aversiveness, is observed in these plots, as well as the ability of our model to predict it.

**Supplementary Figure 8.**
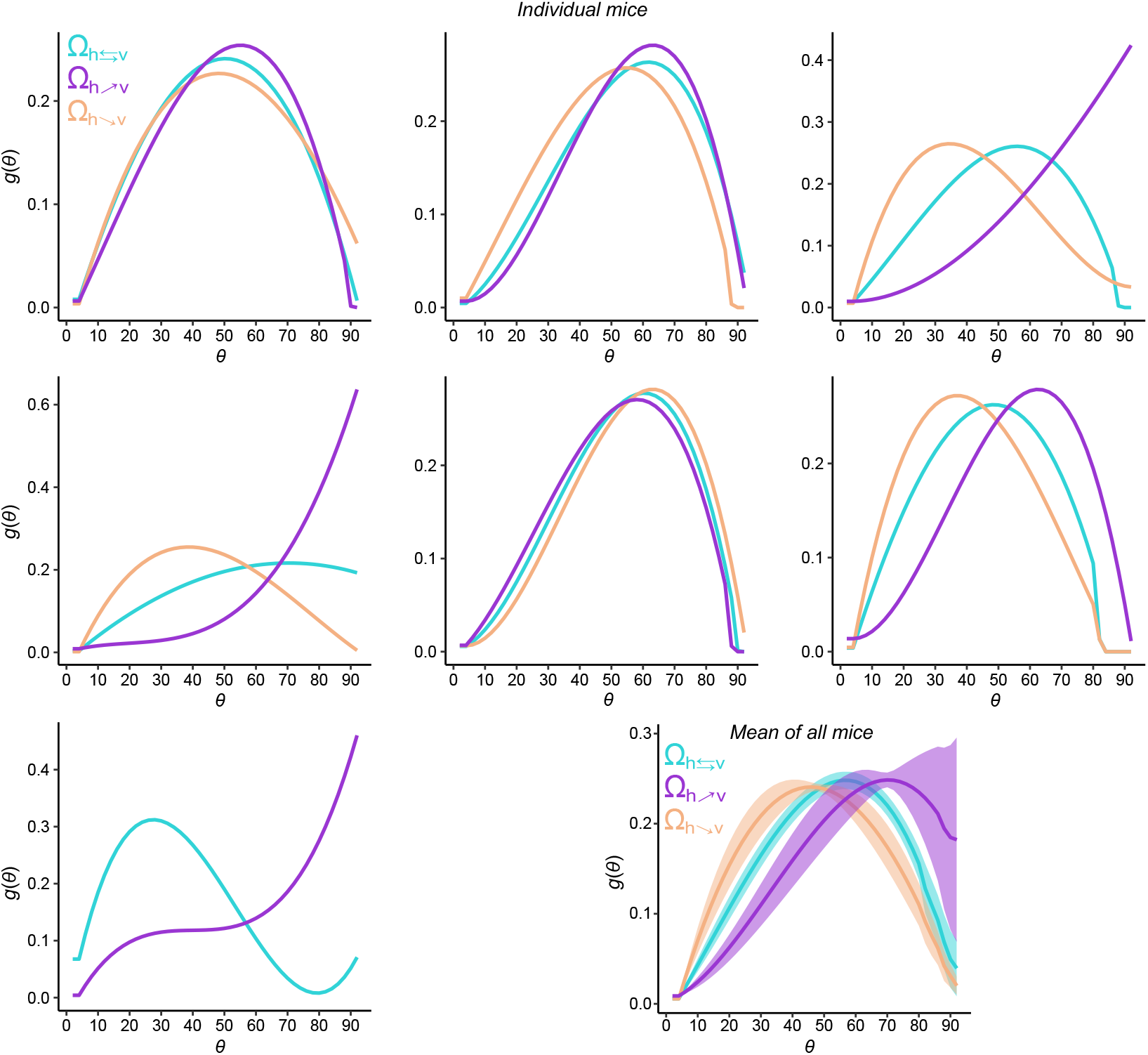
Individual animals show adaptation to different reward environments. Resource allocation functions *g*(*θ*) for each reward environment Ω and each individual mouse. Bottom right plot shows the means of all mice that had all reward environments (top two rows). Shaded areas show standard 95% bootstrap CIs. Individual animals show the trend of increasing the resources allocated areas of stimulus space with higher reward sizes. Gain shifts to more horizontal orientations in 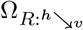, as more reward is dedicated to that portion of stimulus space in this reward condition. The effect is the opposite for 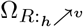, and in between those two is the Ω_*R*:*h*⇄*υ*_, as would be intuitive looking at reward mappings in Figure 1E.

**Supplementary Figure 9.**
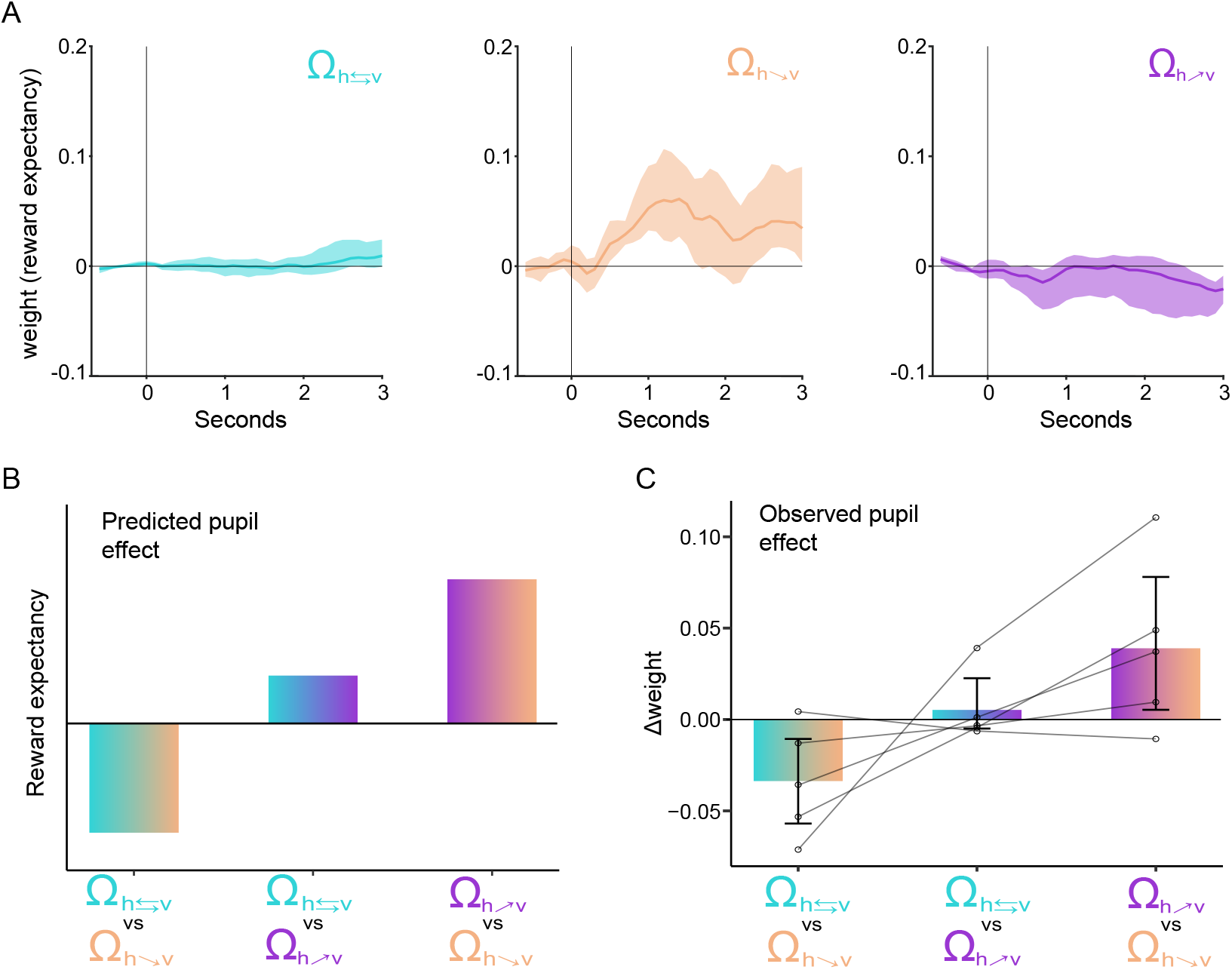
Reward expectancy effect on pupil. (A) Weights of expected reward effect on phasic pupil response over time for each reward environment Ω. Aligned to stimulus onset. Shaded areas show 95% bootstrap CIs. (B) Predicted reward environment pairwise differences in effect of reward expectancy on phasic pupil response. (C) Observed pairwise reward environment differences in weights of expected reward effect on phasic pupil response. Averaged from stimulus onset to 3 seconds post onset. Error bars show 95% CIs and lines show differences in individual mice.

**Supplementary Table 1.** Stimulus-reward mapping. The amounts of milkshake dispensed at a given correct side angle in a given reward environment Ω. In 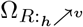, reward size increases from horizontal to vertical. The opposite is true for the 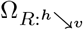 mapping, whereas same size of reward is assigned to all angles in the Ω_*R*:*h*⇄*υ*_ reward mapping condition.

| Reward Mappings | Correct side angle (degrees) |  |  |  |  |  |  |  |  |  |
| --- | --- | --- | --- | --- | --- | --- | --- | --- | --- | --- |
|  | 0 | 10 | 20 | 30 | 40 | 50 | 60 | 70 | 80 | 90 |
| $\Omega_{R:h \nearrow v}, \mu\text{L}$ | 1 | 1.8 | 2.6 | 3.3 | 4.1 | 4.9 | 5.7 | 6.4 | 7.2 | 8 |
| $\Omega_{R:h \searrow v}, \mu\text{L}$ | 8 | 7.2 | 6.4 | 5.7 | 4.9 | 4.1 | 3.3 | 2.6 | 1.8 | 1 |
| $\Omega_{R:h \leftrightarrow v}, \mu\text{L}$ | 5 | 5 | 5 | 5 | 5 | 5 | 5 | 5 | 5 | 5 |

